# Structural basis of internal peptide recognition by PDZ domains using molecular dynamics simulations

**DOI:** 10.1101/575233

**Authors:** Neetu Sain, Debasisa Mohanty

## Abstract

PDZ domains are important peptide recognition modules which usually recognize short C-terminal stretches of their interaction partners, but certain PDZ domains can also recognize internal peptides in the interacting proteins. Due to the scarcity of data on internal peptide recognition and lack of understanding of the mechanistic details of internal peptide recognition, identification of PDZ domains capable of recognizing internal peptides has been a difficult task. Since Par-6 PDZ domain can recognize both C-terminal and internal peptides, we have carried out multiple explicit solvent MD simulations of 1 μs duration on free and peptide bound Par-6 PDZ to decipher mechanistic details of internal peptide recognition. These simulations have been analyzed to identify residues which play a crucial role in internal peptide recognition by PDZ domains. Based on the conservation profile of the identified residues, we have predicted 47 human PDZ domains to be capable of recognizing internal peptides in human. We have also investigated how binding of CDC42 to the CRIB domain adjacent to the Par6 PDZ allosterically modulate the peptide recognition by Par6 PDZ. Our MD simulations on CRIB-Par6_PDZ di-domain in isolation as well as in complex with CDC42, indicate that in absence of CDC42 the adjacent CRIB domain induces open loop conformation of PDZ facilitating internal peptide recognition. On the other hand, upon binding of CDC42 to the CRIB domain, Par6 PDZ adopts closed loop conformation required for recognition of C-terminus peptides. These results provide atomistic details of how binding of interaction partners onto adjacent domains can allosterically regulate substrate binding to PDZ domains. In summary, MD simulations provide novel insights into the modulation of substrate recognition preference of PDZ by specific peptides, adjacent domains and binding of interaction partners at allosteric sites.

## Introduction

PDZ domains are peptide recognition modules which mediate protein-protein interactions during assembly of protein signaling complexes in a variety of biological processes ^1^. PDZ mediated interactions generally have low affinity but high specificity. Disruption of PDZ mediated interactions leads to a variety of human diseases like neurological disorders and cancer ^2–3^. PDZ domains constitute one of the largest family of globular domains in the human proteome and *in silico* analysis by te Velthuis *et al.* has revealed 267 occurrences of the PDZ domains in 154 human proteins ^4^. However, PDZ mediated interaction network of human proteome is yet to be comprehensively characterized. Hence, several studies have attempted to decipher substrate specificity of PDZ domains. PDZ domains usually recognize five to seven residues long peptide stretches present at C-terminus of their interaction partners ^5–6^. These peptide recognition domains are typically ∼80-90 amino acids long and have a conserved fold consisting of 5-6 β-strands and 2-3 α-helical segments. A hydrophobic binding pocket is formed by the residues from the βB sheet and αB helix along with conserved carboxylate-binding loop residues that interact with the free C-terminus of the peptide ligand **(Figure 1A)**. Since peptide binding pocket of PDZ domains are closed at one end by the carboxylate binding loop and harbor conserved residues which can stabilize the free carboxy terminus of interaction partners, this binding pocket geometry is believed to be primary determinant of C-terminus peptide recognition by most PDZ domains **(Figure 1)**^6^.

**Figure 1.**
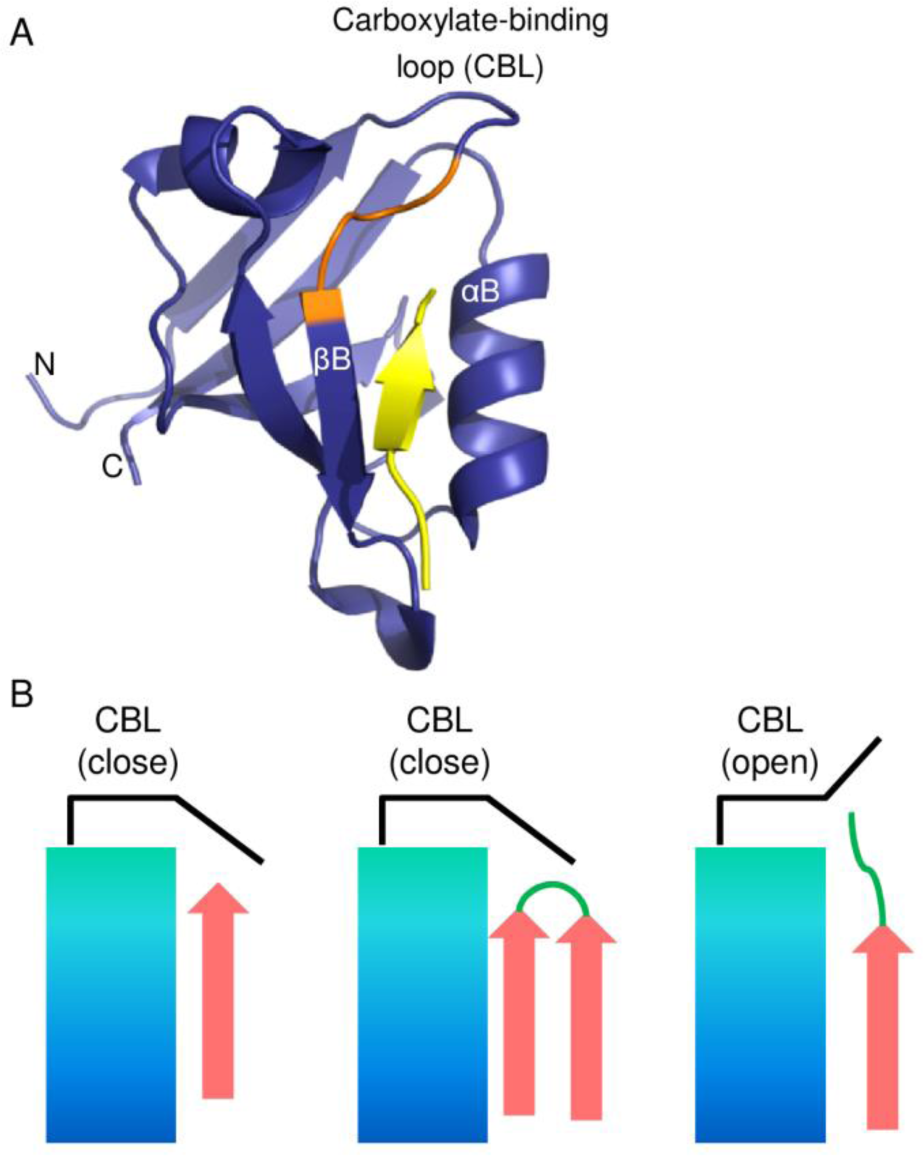
PDZ domain and its binding modes. A) Representative structure of PDZ domain with peptide shown in yellow color. The corresponding conserved GLGF motif of carboxylate binding loop is shown in orange color. B) Cartoon representation depicting different binding modes of PDZ domain with canonical (closed) and non-canonical (open) carboxylate-binding loop conformation.

Although C-terminal peptide recognition is the canonical or most dominant mode of interaction for PDZ domains, there are examples of PDZ domains which can also recognize internal peptide (not present at C-terminus) stretches of their interaction partners. As compared to C-terminal peptide recognition mechanism, our understanding of the mechanistic details of internal peptide recognition by PDZ domains is limited. Hence, majority of the experimental as well as *in silico* studies for deciphering the specificity landscape of PDZ domains have concentrated on C-terminal peptide recognition. However, a recent genome wide screen using random octapeptide yeast two-hybrid library consisting of internal PDZ binding motifs (PBMs) has identified novel 24 PDZ domains ^7^. This suggests that the ability of PDZ domains to bind internal peptides is much more prevalent than previously recognized. Therefore, it is important to understand the mechanism and coverage of internal peptide interactions by PDZ domains so that their interactions can be selectively targeted or modulated by small molecules and peptidomimetics. Internal peptide recognition by PDZ domains require the internal peptide to adopt specific conformations (e.g., β-finger ^8^, cyclic peptides ^9^) which can be accommodated in the closed C-terminal peptide binding pocket or by subtle alteration in the shape and topology of the peptide binding pocket by deforming conformation of the carboxylate-binding loop of the PDZ domain ^10^**(Figure 1B)**. Some of the well-studied PDZ domains whose structures are available in PDB, e.g. Par-6 ^10^, DVL2 ^11^, DLG1-1 and hGIP^12–13^ can recognize both C-terminal and internal peptides by utilizing conformational plasticity of their carboxylate-binding loop. However, it is indeed intriguing how a single PDZ domain can recognize both C-terminus and internal peptide ligands with high specificity by subtle alteration in the conformation of its carboxylate-binding loop. The conformational switch in PDZ domain for C-terminal *vs* internal peptide recognition is also often allosterically regulated as in case of cell polarity protein Par-6.

Par-6 is a multi-domain protein which constitutes a critical component of the cell polarity complex ^14^. It consists of Phox/Bem (PB1) domain, Cdc42/Rac-interactive binding (CRIB) domain and PDZ domain. This single PDZ domain in Par-6 can bind to C-terminal of Crumbs protein as well as an internal peptide stretch in Pals1 protein. However, the binding of Rho GTPase Cdc42 to adjacent 26 amino acid long CRIB domain leads to allosteric transition in CRIB-PDZ resulting in a conformation which binds to C-terminal peptide with a 10 fold higher affinity ^15–16^, while internal peptide can bind to Par-6 PDZ in absence of Cdc42 ^10^. Since crystal structures are available for Cdc42 bound CRIB-PDZ module of Par-6 and Par6 PDZ domain alone in complex with C-terminal as well as internal peptide ligands, analysis of these structures have provided insights into structural basis of C-terminal *vs* internal peptide recognition and allosteric control of substrate recognition by Par-6 PDZ domain. Comparison of the crystal structures of the Par-6 PDZ domain in complex with C-terminal as well as internal peptides revealed that they mainly differ in the conformation of the carboxylate-binding loop ^10^. In the C-terminus peptide complex, the carboxylate binding loop (CBL) is in “closed conformation” and forms a groove to hold the free C-terminus of the ligand peptide. In the internal peptide complex, CBL moves upward to accommodate the residues beyond pseudo C-terminus i.e., forms “open conformation”. It has also been proposed that the salt bridge formation between the lysine present in carboxylate-binding loop and aspartate from internal peptide may be responsible for stabilizing the open conformation. Based on NMR studies in isolated Par-6 PDZ domain (156-255), CRIB-PDZ (130-255), Cdc42 bound CRIB-PDZ and disulfide linked CRIB-PDZ (Q144C/L164C), Whitney *et al.*^16–17^ have proposed that allosteric regulation of conformational transition in Par-6 PDZ domain is mediated by a “dipeptide switch” involving orientation of Leu 164 and Lys 165 residues towards/away from the peptide binding pocket.

Even though the available crystal and NMR structures of Par-6 PDZ domain in complex with different peptide ligands and allosteric effectors have provided extremely valuable structural information to enhance our understanding of the structural details for substrate selection and its allosteric regulation, the role of protein dynamics in selection of C-terminal *vs* internal peptides as well as effect of dynamics on allosteric regulation of substrate binding has not been completely understood. Since, long time scale atomistic simulations can provide unprecedented mechanistic details on interplay between conformational dynamics and binding specificity during molecular recognition processes, simulation studies are often referred as “computational microscopy” for deciphering conformational transitions. Atomistic MD (molecular dynamics) simulations have been extensively used to investigate biomolecular complexes ^18^. Several Recent studies have used MD simulations for understanding dynamics of ligand binding ^19^, dynamics of multi PDZ domains^20^ and allosteric interactions^21–22^ associated with C-terminal peptide recognition by PDZ domains. However, internal peptide recognition has not been studied in detail. The C-terminal peptide simulation studies have helped in identifying residue networks of PDZ domains responsible for long range transmission of conformational rearrangements upon ligand binding ^23–24^. The energetic origin of these long range networks have also been deciphered using residue-residue interaction energy correlations ^25^. Simulation studies by Buchli *et al.* has revealed how rearrangement of water network on surface of PDZ leads to the opening of the binding groove of PDZ domain ^26^. A molecular dynamics study on PSD95-PDZ3 showed the role of residues from β2-β3 loop and C-terminal extra domain helix in enhanced binding affinity for peptides, which cannot be inferred from static crystal structure ^27^. Atomistic simulations have also helped in identification of crucial specificity determining residues of PDZ domain, which make stable contacts with substrate peptide during MD simulations, but are located beyond contact distance in the static structure. Based on MD simulations Steiner *et al.*^28^ have proposed the involvement of “conformational selection” as a possible mechanism for C-terminal peptide recognition by PDZ3 domain of PSD95. A more recent study on CRIB-PDZ involving Markov State Model (MSM) analysis of MD trajectories has investigated the role of allosteric effector on C-terminal peptide recognition ^22^. Even though MD simulation studies on C-terminal peptide recognition by PDZ domain have provided valuable insights which can be obtained from analysis of crystal and NMR structures alone, internal peptide recognition by PDZ domain and allosteric regulation of C-terminal *vs* internal peptide recognition has not been investigated using MD simulation studies. Therefore, in this work we have carried out several microsecond scale molecular dynamics simulations on Par-6 PDZ domain to understand structural basis of internal peptide recognition and its allosteric regulation.

To explore the conformational landscape and understand the allostery involved in Par6 PDZ dual specificity, we performed molecular dynamics (MD) simulations of 1 μs in combination with MM-PBSA (Molecular Mechanics-Poisson–Boltzmann Surface Area) analysis on the multiple Par-6 PDZ complexes. These simulations were examined to identify the residues/features responsible for conformational rearrangements which may further be extrapolated to explore the internal peptide binding capabilities of other PDZ domains. Next to get better insight into recognition mechanism followed by Par-6 PDZ, we simulated both the peptide-bound crystal structures after removing their respective peptide and CRIB-PDZ module with and without CDC42 which led to identification of interaction mediating allosteric communication.

## Results

### Plasticity of carboxylate binding loop of Par6 PDZ

The availability of the crystal structures of C-terminal (PDB entry 1RZX) ^15^ and internal peptide bound complexes of Par-6 PDZ (PDB entry 1X8S) ^10^ allowed us to analyze how the same PDZ domain alters its binding pocket geometry for specific recognition of two distinct ligands. **Figure 2** shows the superposition of the crystal structures of C-terminal peptide (VKESLV) bound Par-6 PDZ domain (1RZX) on internal peptide (HREMAVDCP) bound crystal structure of the same Par-6 PDZ domain (1X8S). As per the numbering scheme followed for PDZ binding peptides in the literature, the residues in the C-terminal peptide VKESLV are −5 to 0 and accordingly the internal peptide is numbered −5 to +3. As can be seen the major portions of PDZ domain in both structures show very good superposition, which conformational differences are restricted to the carboxylate binding loop. The HREMAV segment of the internal peptide superposes well with C-terminal peptide VKESLV. However, while in case of C-terminal peptide the carboxylate binding loop moves towards the C-terminal carboxylate of peptide ligand and adopts closed loop conformation, in case of internal peptide complex carboxylate binding loop moves upward to accommodate the sequence stretch DCP of the internal peptide resulting in open loop conformation. Since major objective of our study was to investigate the dynamics of conformational rearrangements in Par-6 PDZ domain in response to changes in the sequence and structure of the peptide ligand, we set up a series of 1 μs explicit solvent molecular dynamics (MD) simulations on open and closed loop structures of Par-6 PDZ domain in complex with cognate and non-cognate peptide ligands **(Table 1)**. As described in methods section, while structure of cognate complexes corresponded to PDB entries 1RZX and 1X8S were available, non-cognate complexes were generated by exchanging the ligand peptides **(Figure S1)**. We wanted to investigate if the MD simulations on non-cognate complexes will lead to open to closed state conformational transitions or *vice versa* as per the bound ligand. In order to predict the preferred loop conformation of the ligand free Par-6 PDZ domain, 1 μs explicit solvent simulations were carried out for PDZ domain alone starting from open and closed states **(Figure S1)**.

**Table 1.** Details of explicit solvent MD simulations carried out for Par6 PDZ domain.

| S. No. | System | Description | Time |
| --- | --- | --- | --- |
| 1 | Cognate Par6 internal complex (1X8S) | Internal peptide bound to PDZ having open conformation of loop | 1μs |
| 2 | Non-cognate Par6 internal complex | Internal peptide transformed to PDZ having closed conformation of loop | 1μs |
| 3 | Cognate Par6 C-terminal complex (1RZX) | C-terminal peptide bound to PDZ having closed conformation of loop | 1μs |
| 4 | Non-cognate Par6 C- | C-terminal peptide transformed to PDZ | 1μs |
|  | terminal complex | that has open conformation of loop |  |
| 5 | Cognate Par6 internal complex without peptide | Internal peptide coordinates are removed | 1μs |
| 6 | Cognate Par6 C-terminal complex without peptide | C-terminal peptide coordinates are removed | 1μs |
| 7 | CDC42-CRIB-(Par6)PDZ (1N73) | No peptide and has close conformation | 1μs |
| 8 | CRIB-(Par6)PDZ | CDC42 is removed from 1N73 | 1μs |

**Figure 2.**
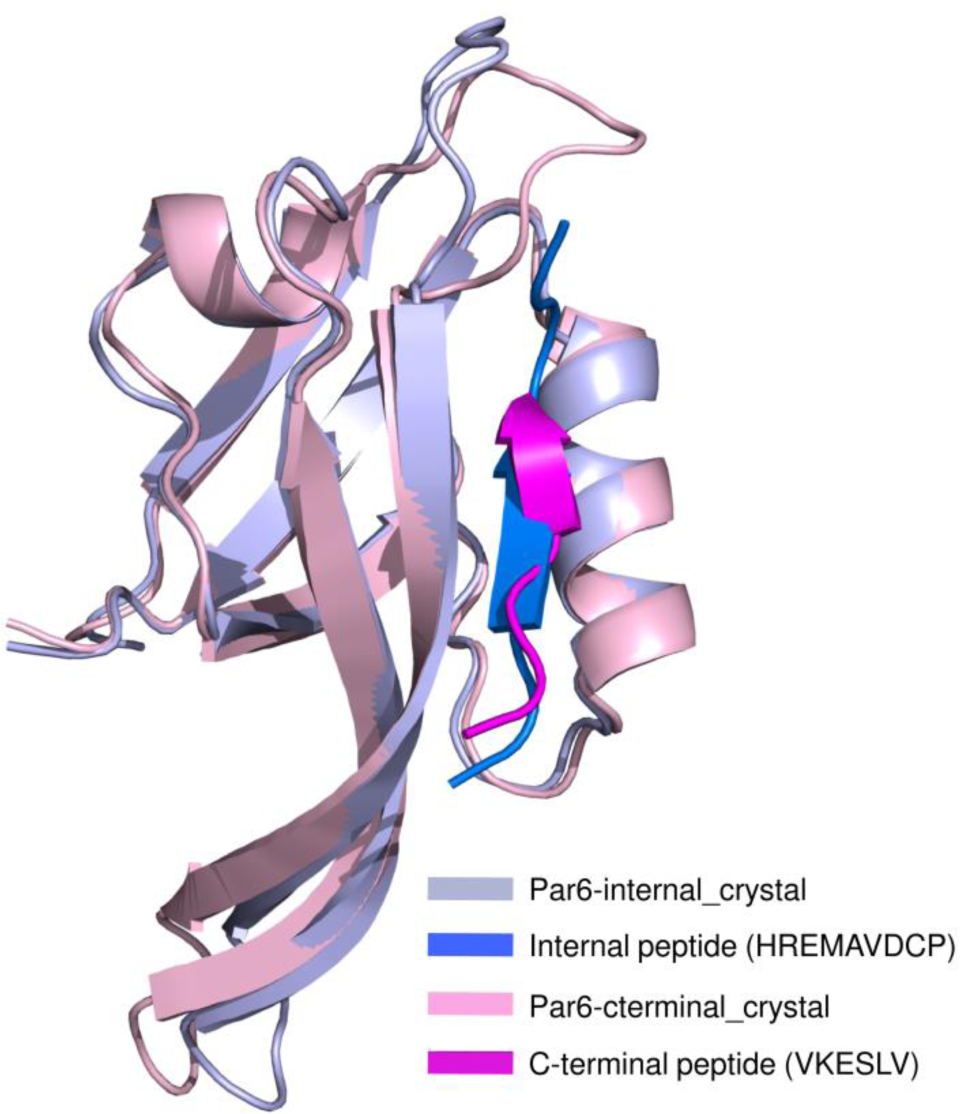
Superimposition of Par6 C-terminal and internal complex. It clearly shows the distinct conformations of carboxylate binding loop depending on the interacting peptide.

In order to facilitate easy analysis of population of various conformational states sampled during the simulations, the conformers in different trajectories were clustered as described in methods section. This resulted in a total of 12 clusters. Representative structures from each of these 12 clusters were compared with closed and open states of Par-6 PDZ seen in crystal structures of c-terminal and internal peptide complexes. **Figure 3** shows the loop RMSD (Root Mean Square Deviation) matrix for these 14 structures, where each box depicts heat map for RMSDs for the loop region when a given pair of structures are optimally superposed using TM-Align software. As can be seen from **Figure 3**, the 12 representative structures from the clusters of conformers sampled during MD simulations can be classified broadly into three groups in terms of their loop RMSD values. While majority of them are similar (loop RMSDs in the range of 0 to 3Å) to the closed loop structure seen in crystal structure of C-terminal peptide bound PDZ domain, three are close (loop RMSDs in the range of 0 to 2Å) to the open loop structure seen in the crystal structure of internal peptide bound PDZ domain. On the other hand, representative structures from two clusters sampled by peptide free PDZ domain constitute the third group which have loop RMSDs from closed loop structures in the range of 3 to 5Å and from the open loop structure in the range of 6 to 7Å. **Figure 4** shows superposition of representatives from each of these three groups along with C-terminal and internal peptide bound Par-6 PDZ crystal structures which were used as reference structures for this analysis.

**Figure 3.**
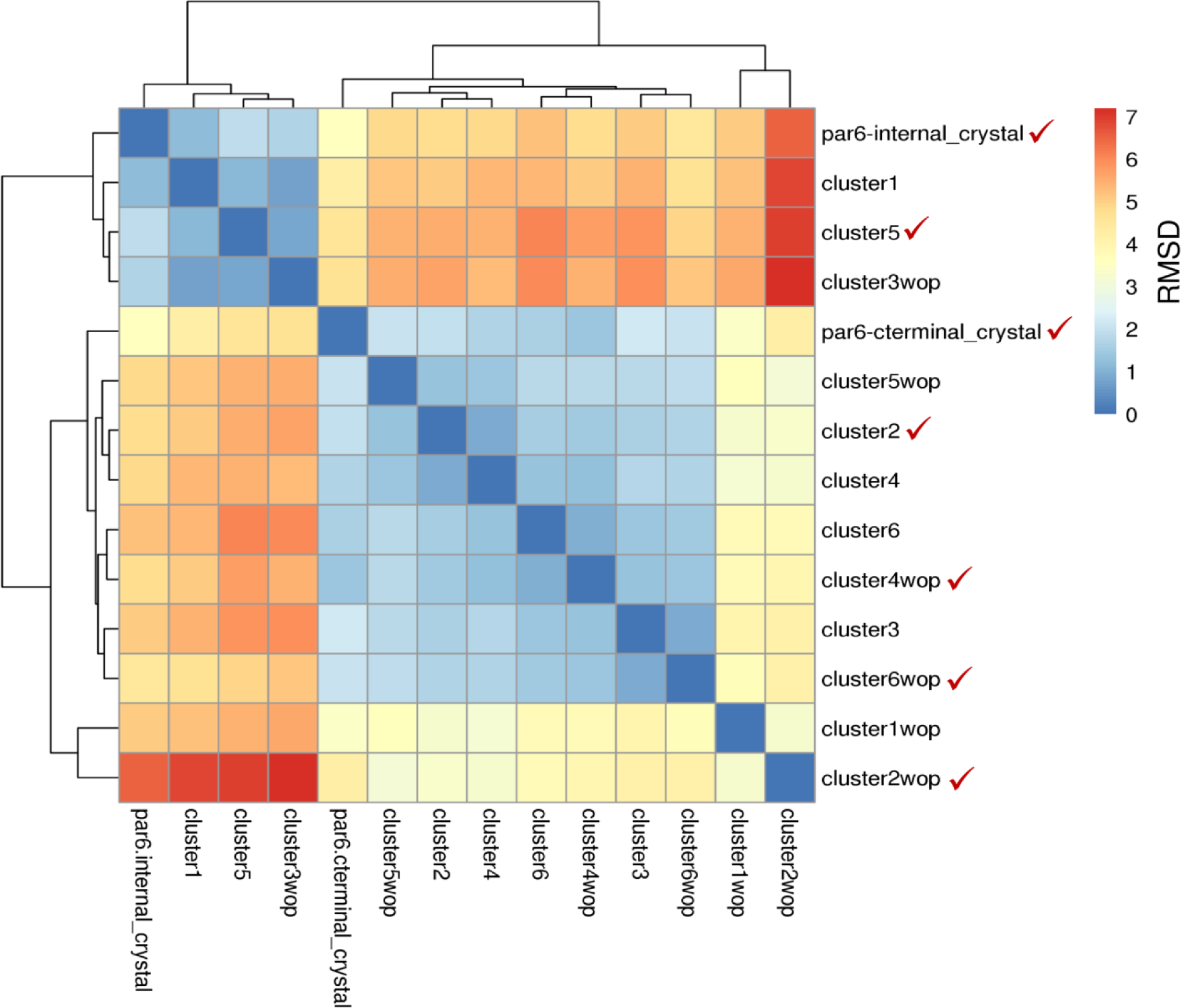
Heatmap showing the hierarchical clustering of representative members based on RMSD of carboxylate binding loop. Representative members were taken from each of the 12 clusters obtained after clustering of ‘cognate & non-cognate’ and ‘cognate & without peptide’ trajectories. In this, crystal structures of C-terminal and internal peptide complexes were also included for reference depicting close and open conformation of loop respectively.

**Figure 4.**
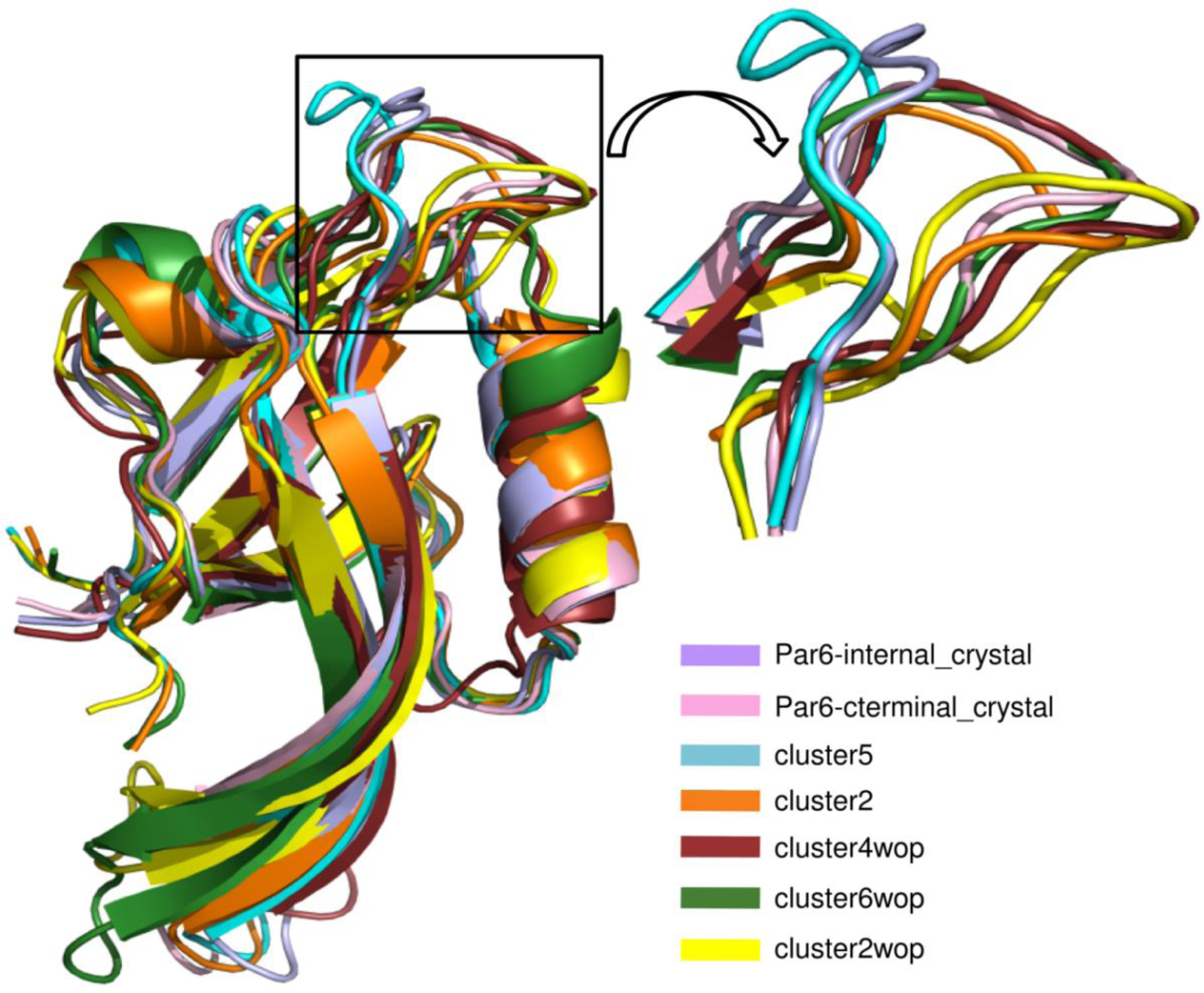
Superimposition of representative structures from the hierarchical clustering depicted in Figure 3. Five representative members marked with tick in Figure 3 were selected and superimposed with structure of C-terminal and internal peptide complex. These structures can be divided into two groups mainly based on conformation of carboxylate binding loop i.e., close and open. Though a representative structure from cluster2wop shown in yellow color is in close conformation but it is distinct from close conformation of cognate C-terminal complex.

Detailed analysis of the simulation results revealed ligand-induced conformational changes in the carboxylate-binding loop of Par-6 PDZ domain. **Figure 5** shows starting structure and the last frame from the trajectory after 1 μs for each of the four peptide bound Par6-PDZ simulations. As can be seen, the starting structure for the non-cognate C-terminal complex had carboxylate binding loop in open conformation and C-terminal peptide in the binding pocket. During the 1 μs of simulation the carboxylate-binding loop of PDZ has moved downwards to form a groove which closes the binding pocket at one end and facilitates favorable interactions of carboxylate binding loop with the free C-terminus of peptide ligand and finally at the end of the simulation reached closed conformation similar to the cognate C-terminal complex **(Figure 5A and 5B)**. In case of non-cognate internal peptide complex, when internal peptide coordinates were transformed onto the Par6-PDZ structure having closed conformation for carboxylate-binding loop, Asp at +1 position of the peptide ligand had steric clashes with Pro171 of the PDZ domain. These clashes were removed after minimization of the system by slight movement in the loop and peptide **(Figure S2)**, but the carboxylate-binding loop in the starting structure used for MD simulation was in closed conformation. However, during the simulation the carboxylate binding loop moved away from the peptide binding groove, but the residues at +1 to +3 position of the internal peptide interacted with the loop in open conformation. At the end of the 1 μs simulation non-cognate internal peptide complex achieved a peptide bound open conformation similar to the cognate internal peptide complex **(Figure 5C and 5D)**.

**Figure 5.**
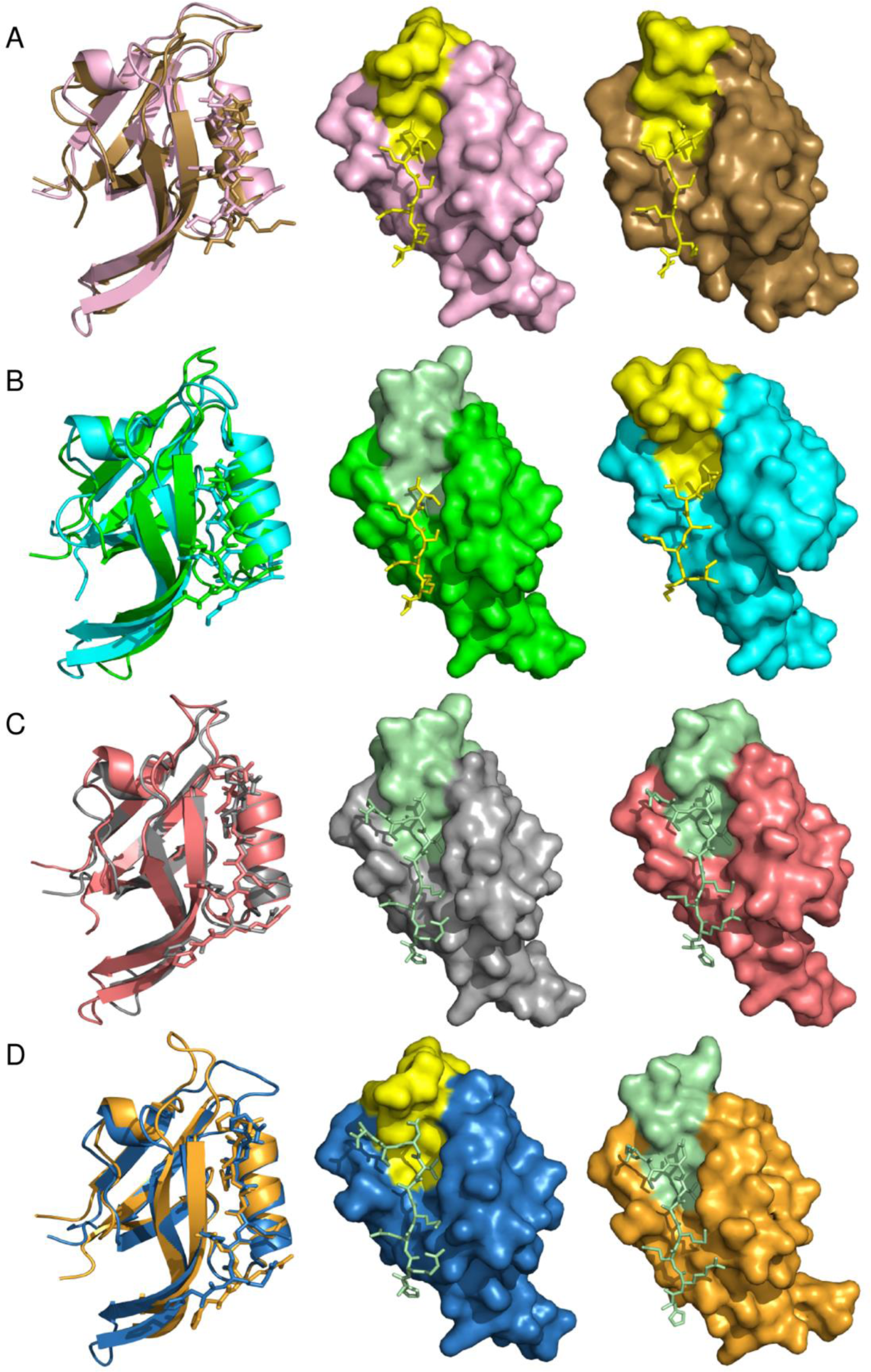
Flexibility of Par-6 PDZ carboxylate-binding loop. Conformations of carboxylate-binding loop in (A) cognate C-terminal complex, (B) non-cognate C-terminal complex, (C) cognate internal complex and (D) non-cognate internal complex for their starting structure and last structure of the trajectory are shown. Middle structure from each figure was used as the starting system for the simulation and right panel is showing the structure obtained at the end of the simulation in surface representation. C-terminal peptide and close conformation of carboxylate-binding loop (residues 8-19) are shown in stick and surface representation respectively in yellow color. Internal peptide and altered i.e., open carboxylate-binding loop (residues 8-19) are represented similarly in pale green color. Left panel shows the superposition of first and the last structure of simulations in the same color as their surface representations.

**Figure S3** shows the RMSF (root mean square fluctuations) plots of cognate and non-cognate complexes plotted separately for C-terminal and internal peptide complex simulations. As expected high fluctuations were observed for residues in the carboxylate-binding loop (residues 9-16) of non-cognate complexes as compared to the cognate structures. Overall, the simulations indicate that carboxylate-binding loop of Par-6 PDZ domain is flexible enough to attain open or closed conformation depending on the type of peptide peptide ligand it binds i.e. C-terminal or internal peptide. These results are indeed encouraging because our MD simulations can reliably reproduce the ligand induced conformational changes observed in experimental studies on Par-6 PDZ domain.

### Ligand induced ‘closed to open’ state conformational transition in Par-6 is faster than ‘open to closed’ state transition

In order to analyze in detail the dynamics of the ligand induced conformational changes in various cognate and non-cognate PDZ-peptide complexes, the RMSDs for the entire trajectories with respect to the initial structures were plotted against simulation time (**Figure 6**). However, as discussed earlier all the conformers sampled in all four PDZ-peptide complexes can be grouped into six distinct clusters which primarily differ in terms of orientation of the carboxylate binding loop. Therefore, in order to decipher the loop movement during the simulation, each time point in the RMSD *vs* time plot (**Figure 6**) was colored as per the cluster to which the corresponding structure belongs. For each trajectory **Figure 6** also shows the superposition of the representative structures of the clusters sampled during the simulation along with the reference crystal structures of closed and open loop conformations of Par-6 PDZ domain. Percentage occupancy of individual clusters for each MD run is given in **Table 2**. **Figure 6A** shows the superposition of representative members from five of the clusters which are populated by the C-terminal cognate complex conformers, while **Figure 6B** shows time duration for which each of these five clusters were sampled during the 1 µs trajectory. As can be seen from **Figure 6B** and **Table 2**, percentage occupancy of cluster 4, cluster 2 and cluster 3 are 51.9%, 28.7% and 14% respectively, while cluster 6 and cluster 5 have percentage occupancies of 5.3% and 0.1% respectively. Since clusters 2, 3, 4 and 6 correspond to the closed conformation of the carboxylate-binding loop (**Figure 3 and 4)**, the cognate C-terminal complex mostly sampled closed loop conformations throughout the trajectory, while the open loop conformation corresponding to cluster 5 (**Figure 4**) was sampled for lesser time during 1 µs MD run (**Figure 6A, 6B and Table 2**).

**Table 2.**
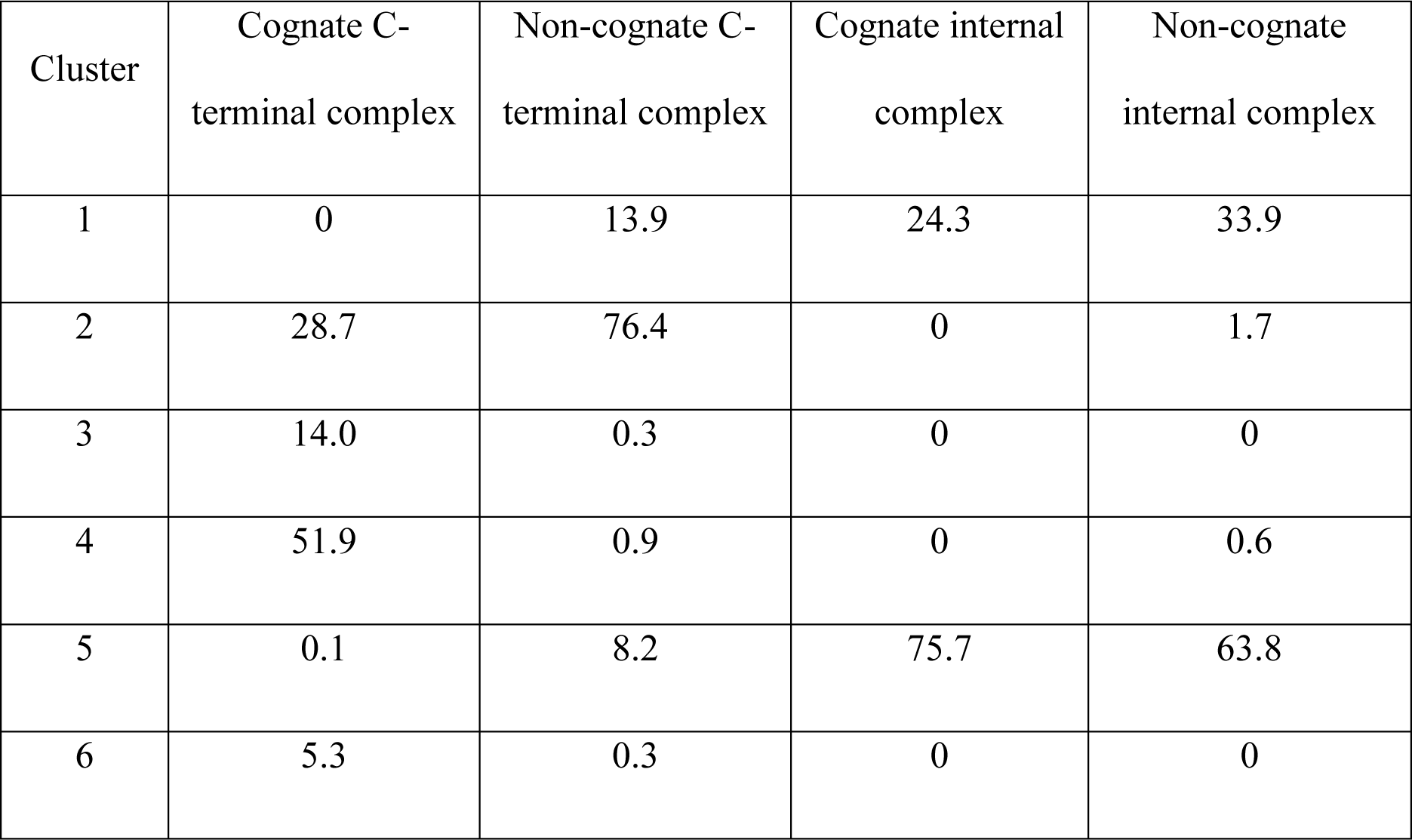
Percentage occupancy of the clusters formed by the trajectories for four simulations on Par-6 PDZ domain in complex with cognate and non-cognate peptide substrates. (Snapshots from all four simulations were clustered together).

**Figure 6.**
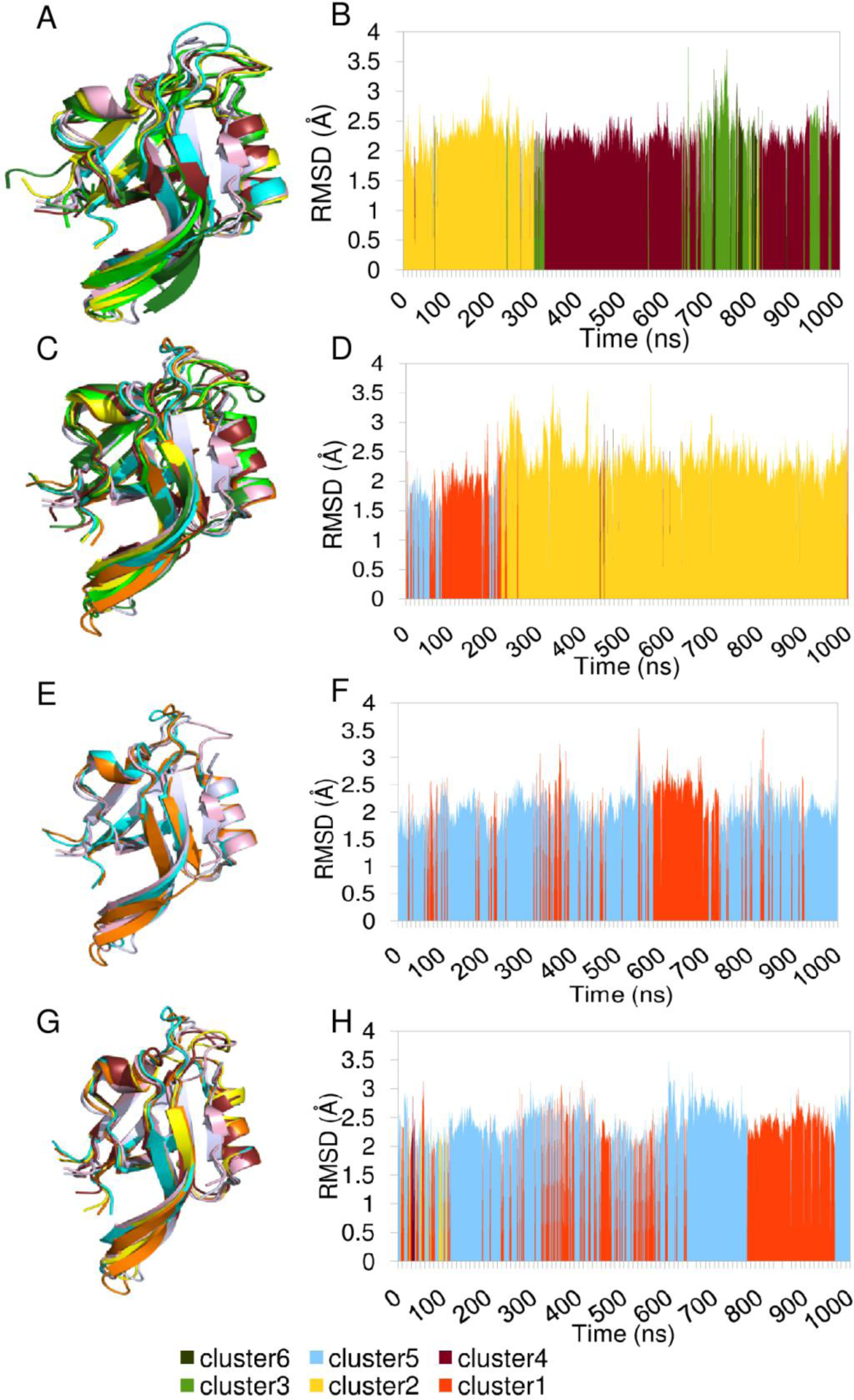
Cluster analysis of cognate and non-cognate C-terminal and internal Par6 PDZ complex trajectories. Cluster analysis was done using kClust on the basis of RMSD. (A and C) superimposition of representative members taken from each cluster of cognate and non-cognate C-terminal complex conformers. (B and D) RMSD vs time plot for cognate and non-cognate C-terminal complex which are colored cluster-wise. (E and G) are showing the overlaid representative members of cognate and non-cognate internal complex clusters. (F and H) are RMSD vs time plots for cognate and non-cognate internal complex. Different colors represent different clusters and with the same color its representative member is shown in the left panel. Same color coding is followed for all the figures. C-terminal (pink) and internal peptide complex (purple) crystal structures are also shown in overlaid structures for reference to open and close conformation.

In case of non-cognate C-terminal complex (**Figures 6C and 6D**), where the carboxylate-binding loop was in open conformation at the beginning of the MD run, open state conformations corresponding to clusters 1 and 5 (depicted in orange and cyan color) were sampled during first 215 ns of the trajectory with percentage occupancies of 13.9% and 8.2% respectively (**Table 2**). However, after ∼215 ns the carboxylate binding loop moved downwards to interact with the free C-terminus of the peptide ligand and cluster 2 (depicted in yellow) which is very similar to the initial structure of cognate C-terminal complex was sampled for most part of the remaining 785 ns of the simulation with a percentage occupancy of 76.4% (**Figure 6C**, **6D and Table 2**). Other closed loop states corresponding to clusters 3, 4 and 6 (**Figure 3, 4 and 6C**) had populations of only 0.3%, 0.9% and 0.3% respectively (**Table 2** and **Figure 6D**). Thus our simulations revealed that the time scale for open to close state conformational transition was around 215 ns.

The 1 µs trajectory for the cognate internal peptide complex revealed that, only open states corresponding to clusters 5 and 1 were sampled when internal peptide was bound (**Figure 3, 6E and 6F**). Cluster 5 which is very similar to the crystal structure of open sate had percentage occupancy of 75.7%, while the other open state (cluster 1) had a population of only 24.3% (**Table 2**) and frequent transitions between these two open states were observed. The absence of closed states in the MD trajectory of cognate internal peptide complex can be explained by the fact that, closed conformation of the carboxylate binding loop results in steric clash with Asp at +1 position of internal peptide as mentioned earlier. The non-cognate internal peptide complex was initially in the closed loop conformation of the carboxylate-binding loop and closed loop conformations corresponding to clusters 2 and 4 (yellow and brown color respectively) were visited for only about 23 ns (occupancy of clusters 2 and 4 were 1.7% and 0.6% respectively) during the first ∼110 ns of the 1 µs MD run (**Table 2, Figure 6G and 6H**). During the initial 110 ns of the simulation, the carboxylate-binding loop showed frequent transition between closed and open states, but open conformation of the carboxylate-binding loop was stabilized after 120 ns and during the remaining 890 ns of the simulation only open states corresponding to the clusters 5 (population 63.8%) and 1 (population 33.9%) were sampled. At the end of the 1µs run non-cognate internal complex converged to cluster 5 which was very similar to the crystal structure of the open state. Thus our simulations revealed that the ‘open to closed’ state conformational transition for Par-6 PDZ domain induced by C-terminal peptide binding was a slower process compared to ‘closed to open’ state conformational change induced by C-terminal peptide binding.

### Ligand free Par-6 PDZ domain prefers closed loop conformation

In order to identify the preferred conformation of Par-6 PDZ domain in isolation (i.e. without bound peptide or N-terminal CRIB domain), we removed the coordinates of bound peptides from crystal structures of cognate C-terminal and cognate internal peptide complex which corresponded to closed and open state respectively. Explicit solvent MD simulations were performed for 1 μs each starting from ligand free closed and open states (**Table 1**). As can be seen from the RMSF (root mean square fluctuations)plots for these two simulations **(Figure S4 and S5)**, there are very significant movements in the carboxylate binding loop region indicating large scale conformational rearrangements between the closed and open states. The simulations starting from the open conformation corresponding to the internal peptide complex showed higher degree of conformational changes. Trajectories for both the ligand free simulations were clustered along with trajectories for cognate C-terminal and cognate internal peptide complexes to investigate if there were overlaps between conformations sampled in ligand bound and ligand free Par-6 PDZ domains. As mentioned earlier the conformers in these four trajectories formed six distinct clusters, namely cluster1wop to cluster6wop and **Figure 3** shows comparison of the structural similarities of the representative structures from these six clusters with the representative structures from clusters sampled by simulations on cognate and non-cognate complexes. As can be seen from **Figure 3** cluster1wop and cluster2wop are similar to each other in terms of loop conformation and they also have higher RMSDs from the loops in both closed and open states. Cluster5wop is most similar to the crystal structure of closed state, while cluster3wop is closer to the crystal structure of the open state. On the other hand cluster4wop and cluster6wop are closer to the closed state structure. **Figure S6** shows superimposition of representative structures from each of the clusters sampled during the trajectory as well as RMSD vs time plot along with depiction of the cluster to which a given snapshot in the trajectory belongs. **Table S1** shows percentage occupancy of various clusters based on the fraction of simulation time for which given cluster is sampled. As can be seen from **Figure S6B**, the cognate C-terminal complex samples closed state conformations in clusters cluster4wop, cluster5wop and cluster6wop with percentage occupancies of 50.8%, 32.6% and 1.8% respectively, while population for the open state like clutser3wop is only 0.2%. The clusters cluster1wop and cluster2wop which are more similar to closed state crystal structure (RMSDs 3 to 5Å) than open state crystal structure (RMSDs 6 to 7Å) have populations of 13.8% and 0,8% respectively. On the other hand, the cognate internal peptide complex samples the open state cluster3wop for 99.3% of the simulation time, while all other clusters have populations of less than 1% only (**Figure S6D**). In case of ligand free Par-6 PDZ simulation starting from closed state conformation, cluster1wop has percentage occupancy of 86%, while closed state clusters cluster5wop and cluster6wop have occupancies of 8.2% and 5.2% respectively and open state cluster cluster3wop has occupancy of 0.6% (**Figure S6A**). On the other hand, in case of ligand free Par-6 PDZ simulation starting from open state conformation, cluster2wop has percentage occupancy of 64.8%, while closed state clusters cluster5wop and cluster6wop have occupancies of 17.5% and 12.9% respectively and open state cluster cluster3wop has occupancy of 4.7% (**Figure S6C**). As can be seen from **Figure 3** cluster1wop and cluster2wop which are maximally populated in both ligand free simulations have RMSDs between themselves in the range of 2 to 3Å and are more similar to closed state structures than open state structures. These results indicate that unbound structure of Par-6 PDZ which started with open conformation predominantly samples an alternate closed loop conformation with a population of 64.8%, while canonical closed and open conformations as in peptide bound PDZ have populations of 30.4% and 4.7% respectively **(Table S1)**.

### Identification of critical residues involved in internal peptide recognition

The various simulations on ligand free as well as ligand bound Par-6 PDZ domains revealed that, in absence of peptide ligand Par-6 PDZ prefers a closed loop conformation and upon binding of C-terminal peptide there is a minor conformational shift within closed loop states. However, upon binding of internal peptide transition to the open loop conformation takes place which is very fast. In an earlier study Perkert *et al*. ^10^ had suggested that the altered conformation of carboxylate bindng loop may be stabilized by salt bridge interaction between the conserved lysine (Lys 165) in the loop and aspartate at +1 position of the internal peptide. They also confirmed the important role of aspartic acid at P(+1) site in internal peptide binding using alanine scanning experiments. We analyzed the distance between the NZ of Lys165 in the carboxylate-binding loop and OD1/OD2 of the Asp+1 of peptide over the simulation trajectories to investigate if our simulation studies agree with the experimental results of Perkert *et al*. As can be seen from **Figure 7A** in case of cognate internal complex (black line) the salt bridge remains stable for most part of 1 µs trajectory. In case of non-cognate internal complex the distance between Lys 165 and Asp at P(+1) is higher during first 150 ns, but after 150ns the salt bridge is formed and remains stable in the remaining part of the trajectory. Thus formation of salt bridge is strongly correlated with the closed to open state conformational transition as seen in cluster analysis plots **(Figure 6H)**.

**Figure 7.**
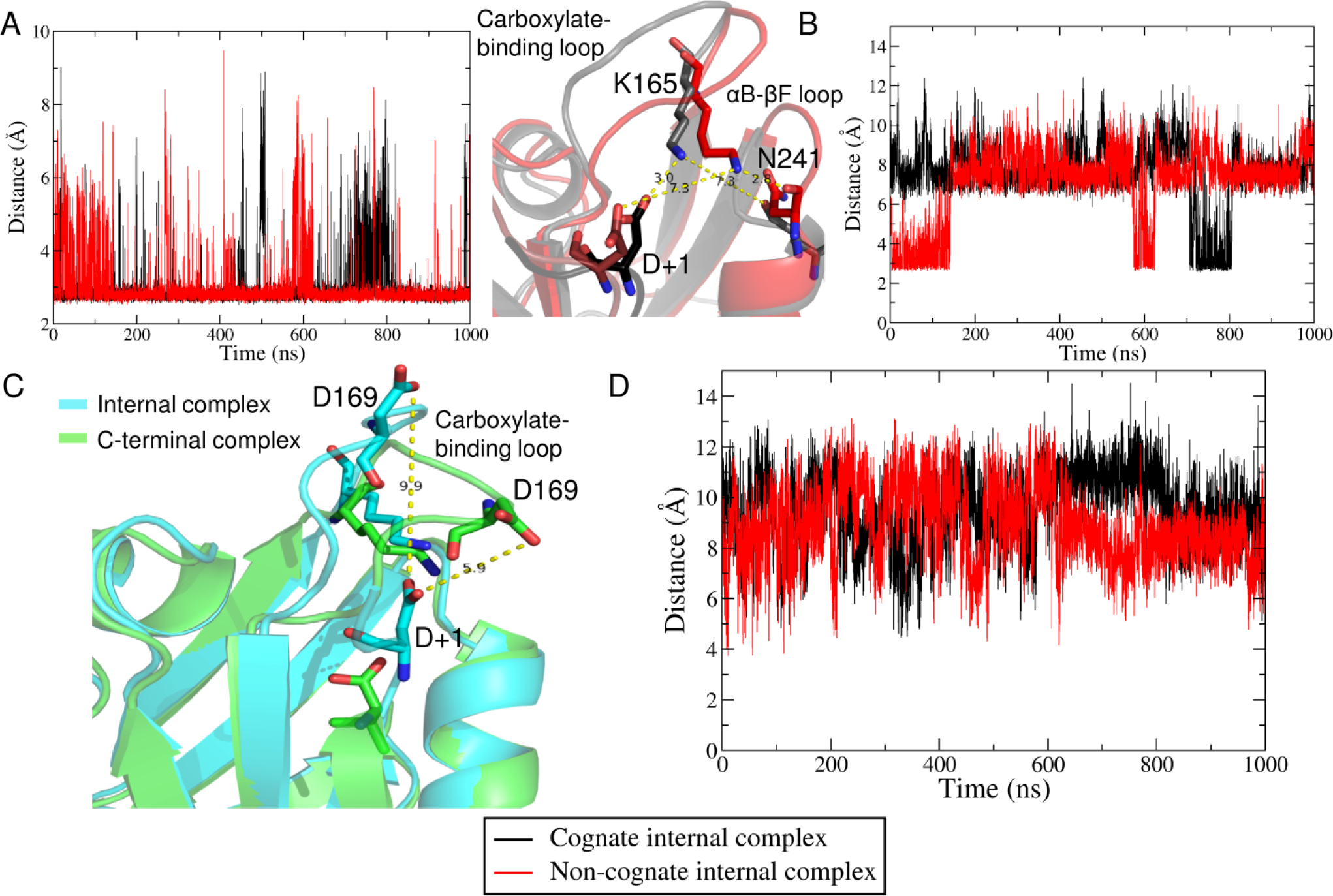
Critical interactions of Par6-PDZ internal complex. (A) Left panel shows the distance plot for Lys165 NZ-Asp+1 OD1/OD2 and right panel shows superimposed cognate (black) and non-cognate (red) internal peptide complexes taken from very start of the simulation with labeled distances between Lys165, Asp+1 and Asn241. (B) Distance plot for Lys165 NZ-Asn241 O, (C) Superimposed Par-6 PDZ C-terminal and internal peptide crystal structures showing orientation and position of Asp169 present in carboxylate-binding loop with respect to the peptide Asp+1, (D) Distance plot for Asp169 OD1/OD2-Asp+1 OD1/OD2.

Even though internal peptide bound conformation was stabilized by Lys 165 to Asp P(+1) salt bridge, the trajectories for both the cognate and non-cognate internal peptide complexes sampled conformations where the distance between Lys-Asp pair considerably increased as can be seen at 600 ns in case of non-cognate complex and 700 to 800 ns in case of the cognate internal peptide complex. Detailed analysis of the representative structures from the trajectories showed the movement in the αB-βF loop of PDZ also. Hence the interactions between residues of carboxylate binding loop and αB-βF loop were analyzed **(Figure S7).** It revealed that when Lys 165:Asp P(+1) salt bridge is broken, amino group of Lys 165 can form hydrogen bonds with backbone carbonyl of Asn 241 from αB-βF loop. **Figure 7B** shows plots for the Lys 165:Asn 241 salt bridge for simulations on non-cognate and cognate internal peptide complexes. As can be seen formation of Lys 165:Asn 241 salt bridge prevents the formation of salt bridge between peptide and carboxylate binding loop, or in other words formation of stable open conformation. Although this analysis indicates the importance of the ion pair formation between carboxylate binding loop and peptide in stabilizing the open conformation, it does not explain why closed conformation of carboxylate binding loop is unfavourable for the binding of internal peptide.

A careful comparison of the side-chain positions and orientations in the carboxylate binding loop of internal and C-terminal peptide crystal structures highlighted the difference in orientation of aspartic acid present at position 169 which is at proximal distance to the Asp+1 residue of peptide in the non-cognate internal peptide complex (**Figure 7C**). **Table S2** shows the distances of this Asp169 of carboxylate binding loop from the Asp P(+1) of internal peptide and Val0 of C-terminal peptide in the starting structures of all four complexes. These distances indicate that repulsion between two aspartates of carboxylate binding loop and peptide in non-cognate internal complex as a drive for close to open state conformational transition. **Figure 7D** shows the plots for Asp 169:Asp (P+1) carbonyl distances throughout the trajectory for the non-cognate and cognate internal peptide complex simulations. Based on these simulation results we propose the following mechanism for internal peptide recognition by Par-6 PDZ domain. The repulsion between Asp P(+1) and Asp 169 of the carboxylate binding loop induces open conformation, but intra molecular interactions in PDZ domain between Lys 165 in carboxylate binding loop and Asn 241 in the αB-βF loop favours closed conformation. Hence during the initial 150 ns of the non-cognate internal peptide complex simulation carboxylate binding loop moves back and forth between open and close states till stable salt bridge interactions between Lys165 and Asp P(+1) is formed and open conformation of loop is stabilized. This repulsion between two aspartates could also be the reason for fast transition from open to closed state in case of non-cognate internal complex.

### Identification of internal peptide recognizing PDZ domains

Next, we examined sequences of the PDZ domains reported to interact with internal peptides in the literature listed by Mu *et al.*^7^ in their study. For efficient alignment, prokaryotic-type or circularly-permuted PDZ domains were kept out of alignment. 10 out of 19 (including Par6) indeed possess negatively charged residue (D/E) in their carboxylate binding loop at corresponding or nearby position (+1/-1) of Asp169 of Par6 as required according to the proposed recognition mechanism **(Figure 8A)**. These all 10 domains also have 1/2/3 residues upstream to aspartate, a positively charged residue in their carboxylate-binding loop. Since Par6 PDZ domain contains two-residue insertion in their carboxylate-binding loop^16^, they could be position of Arg165 of Par6 structurally. These domains may exploit the same internal peptide recognition mechanism as followed by Par-6 PDZ domain. There are other PDZ domains which do not have expected residues and they may be following some other mechanism to interact with internal peptides like forming β-hairpin loop of peptides ^8^ etc. We have found 47 human PDZ domains with aspartate at the corresponding position of Asp169 of Par6 PDZ or at +1/ −1 position with arginine in the carboxylate binding loop when all human PDZ domains were aligned with Par6 PDZ **(Figure S8)** and hence predicted to interact with internal peptides using proposed recognition mechanism.

**Figure 8.**
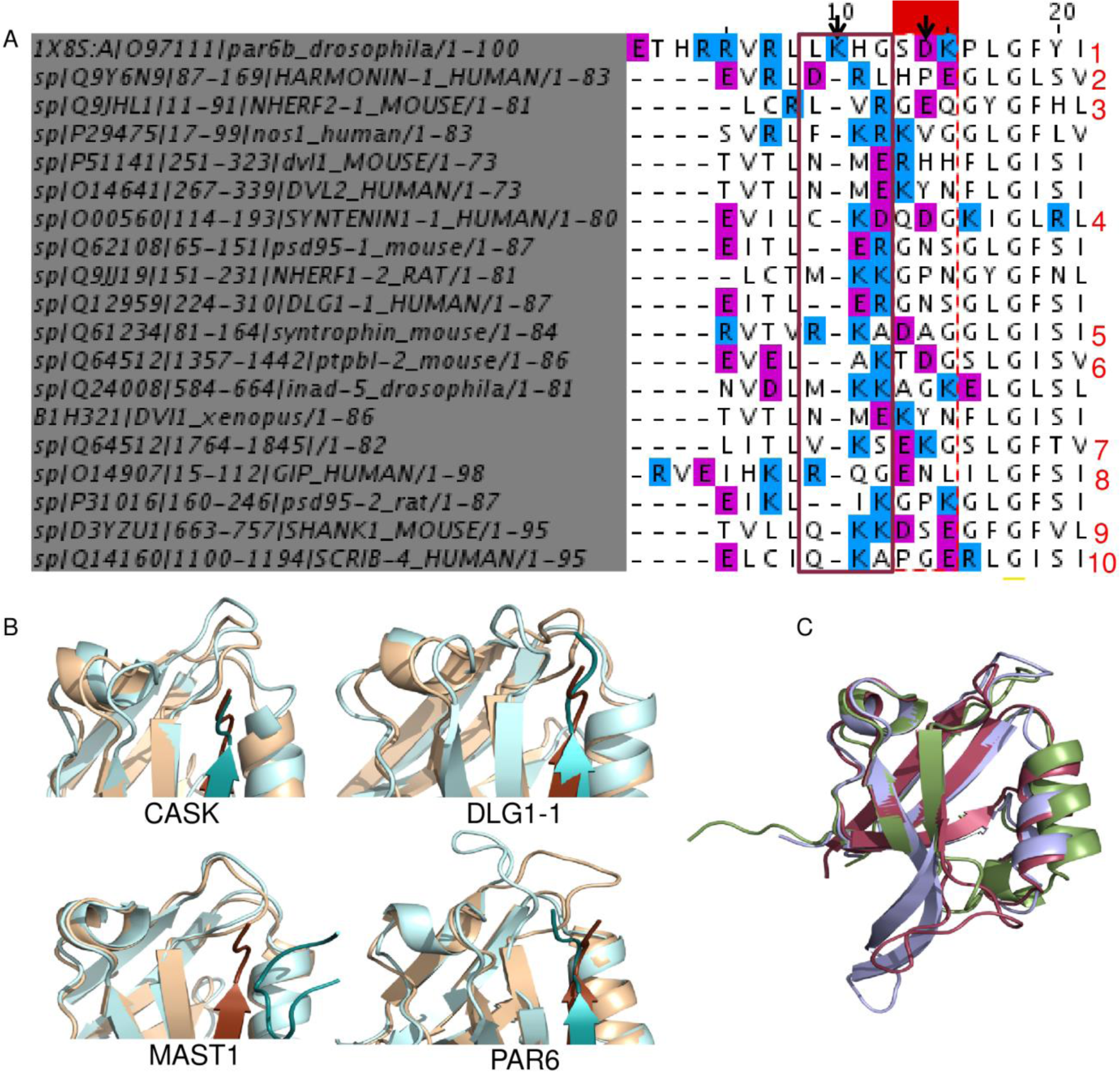
Identification and validation of internal peptide binding PDZ domains. (A) Multiple sequence alignment of internal peptide binding PDZ domains taken from Mu *et al.*^14^. Positively charged residues are colored blue and negatively charged are colored as magenta. Black solid arrows are highlighting the positions of R165 and D169 in Par6 PDZ (PDB ID: 1X8S) numbered as 10 and 15 respectively in alignment. Aspartate present at +1 and −1 position were also considered as positive hit (red dashed box) and Lysine can be seen at one, two or three residue upstream to aspartate (brown solid box). (B) Superimposition of first (brown) and the last structure (blue) of 500ns simulation for CASK, DLG1-1, MAST1 and Par6 (C) Overlaid crystal structures of DLG1-1, CASK and Par6 bound with C-terminal peptide are shown in red, green and purple colors respectively. Peptide is not shown for clarity.

To further validate, we have performed 500 ns MD simulation on CASK PDZ (one of those 47 PDZ domains which have aspartate in carboxylate-binding loop), DLG1-1 PDZ (reported to interact with internal peptide in literature but aspartate in carboxylate-binding loop is not present) and MAST1 PDZ (neither interact with internal peptide nor have aspartate in carboxylate-binding loop) with internal peptide of Par6 PDZ i.e., HREMAVDCP **(Table 3)**. All these PDZ domains have closed carboxylate-binding loop conformation in their crystal structure and HREMAVDCP was transformed on them. As this peptide is not their physiological ligand, a simulation with HREMAV as C-terminal peptide is also run for each of these PDZ domains and average binding energy is calculated using MM-PBSA for last 100ns of trajectory. **Table 3** shows comparable binding energies for C-terminal peptide with all domains but with internal peptide CASK has higher binding affinity than DLG1-1 and MAST1 which is comparable to Par6. Results of simulations are summarized in **Figure 8B** showing superposition of first and last structure of trajectory for all three and Par6 PDZ domain. We can see CASK PDZ is able to accommodate internal peptide with open carboxylate-binding loop, DLG1-1 carboxylate-binding loop is in open conformation but orientation of peptide is different and internal peptide has moved out from the binding pocket of MAST1. It confirms PDZ domains with Asp in their carboxylate-binding loop can accommodate internal peptides having aspartate at +1 position of peptide. Our simulations also showed that the orientation of internal peptide is different in DLG1-1 PDZ which does not have aspartate in carboxylate-binding loop as compared to Par6 PDZ which follows different mechanism for internal peptide recognition. This orientation of peptide has also been observed in a previous report ^29^ for modeled complex of DLG1-1 PDZ where peptide was having additional one residue as compared to C-terminal peptide accommodated by flexible carboxylate-binding loop of DLG1-1 PDZ, and our simulation is suggesting that it can accommodate more residues using same binding mode. Superimposition of Par6 and DLG1-1 PDZ domains **(Figure 8C)** shows DLG1-1 carboxylate binding loop is not in perfectly closed state with C-terminal peptide and it is its inherent flexibility which allows interaction with internal peptide.

**Table 3.**
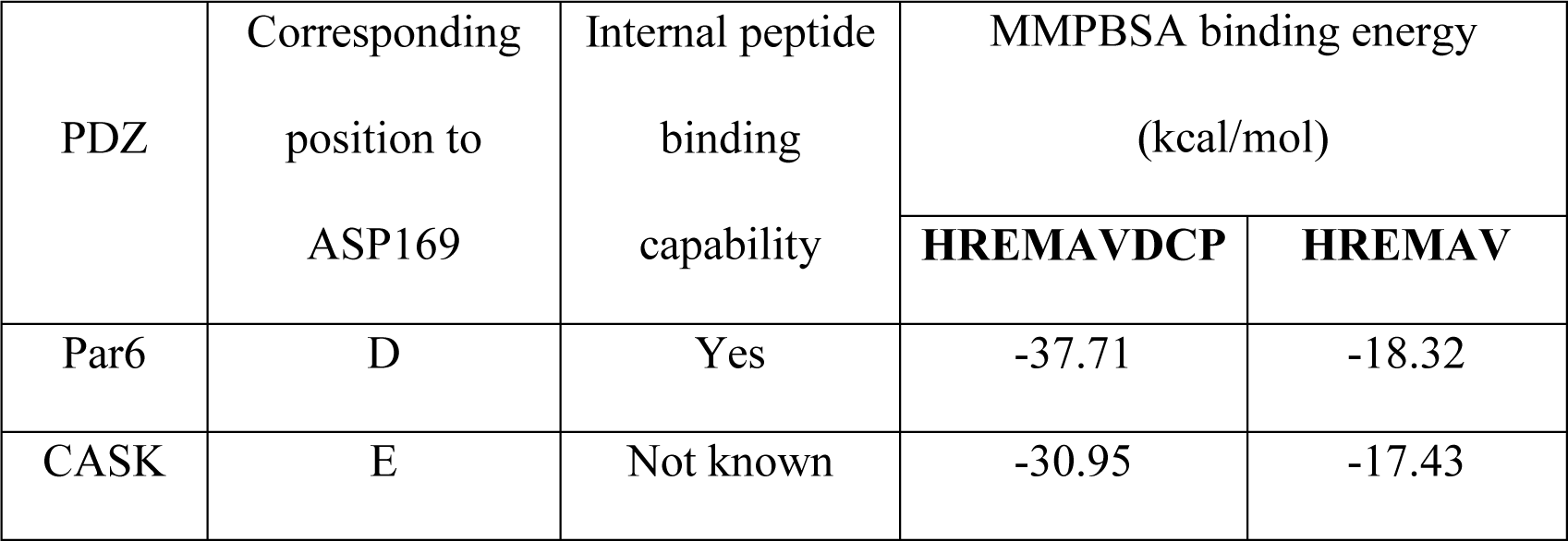
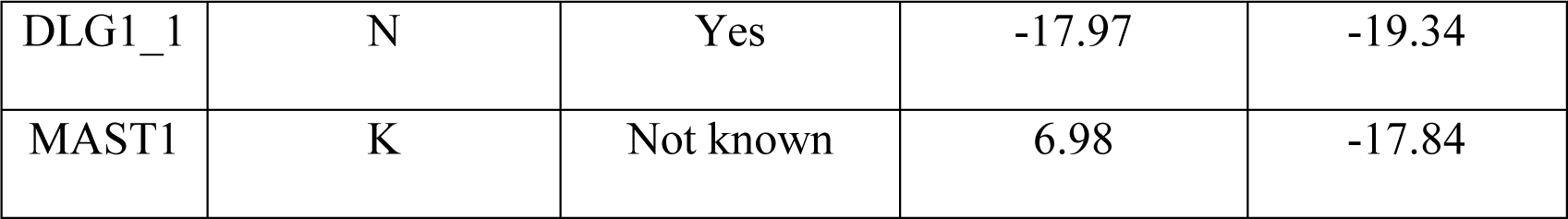
Description of simulations and MM-PBSA energy calculations for CASK, DLG1-1 and MAST1 PDZ for C-terminal (HREMAV) and internal peptide (HREMAVDCP). Par6 PDZ is shown for comparison.

### Calculation of binding free energies using MM-PBSA

We wanted to investigate if preference of Par-6 PDZ domain for internal *vs* C-terminal peptides can be predicted based on their MM-PBSA binding free energy values. MM-PB/SA binding free energy values were computed for cognate as well as non-cognate complexes of Par-6 PDZ using snapshots from the last 500 ns of the 1µs MD trajectories (**Figure S9**). The MM-PBSA binding free energy values were compared with the experimental dissociation constants ^10, 16^ (**Figure S9,** right panel).

The binding energy is higher for the non-cognate C-terminal complex than cognate C-terminal complex while the binding energy for the non-cognate internal complex is comparable to cognate internal complex. We can see from **Figure S9** that the native internal and C-terminal complexes have similar Kd, if CDC42 is bound to CRIB domain adjacent to PDZ domain but without CDC42, it has high dissociation constant. This can explain the high MM-PBSA binding energy for cognate and non-cognate C-terminal complex because the transition from open to close conformation is time taking and less favorable without the help of CDC42 and it is even higher for non-cognate C-terminal complex. It shows MM-PBSA analysis can accurately predict internal versus C-terminal peptide recognition for PDZ domains.

### Effect of CDC42 and CRIB domain on interactions of Par6 PDZ

To examine the role of CDC42 and adjacent domain CRIB in conformational selection of Par6 PDZ **(Figure S10)**, we have simulated complex structure of CDC42 and CRIB-PDZ (PDB ID: 1NF3) and also after removing CDC42 from CRIB-PDZ module for 1µs. As expected, CRIB domain (residue 1-20) is more ordered when bound to CDC42 indicated by its RMSF plot shown in red color in **Figure S11**. The cluster analysis taking both trajectories together was performed as mentioned in methods and they found to be grouped in 8 Clusters. Their percentage occupancy is given in **Table S3**. **Figure 9A** is showing that in presence of CDC42, PDZ domain of Par6 remains in closed loop conformation and goes to only two clusters (lime and orange color). When we removed CDC42, it started in closed conformation of loop (orange) and then transit through intermediate stages of closed and open but finally after 420ns it stably takes up open loop conformation (cyan and brown color) **(Figure 9B)**. It made us to conclude that Par6 PDZ is designed to interact with internal peptide with open loop conformation and only when CDC42 is bound to CRIB, it can interact with C-terminal ligands with higher affinity because CDC42 stabilize the closed conformation of loop favorable for interaction of C-terminal ligands. In this way, PDZ domain is able to have regulated C-terminal peptide interaction.

**Figure 9.**
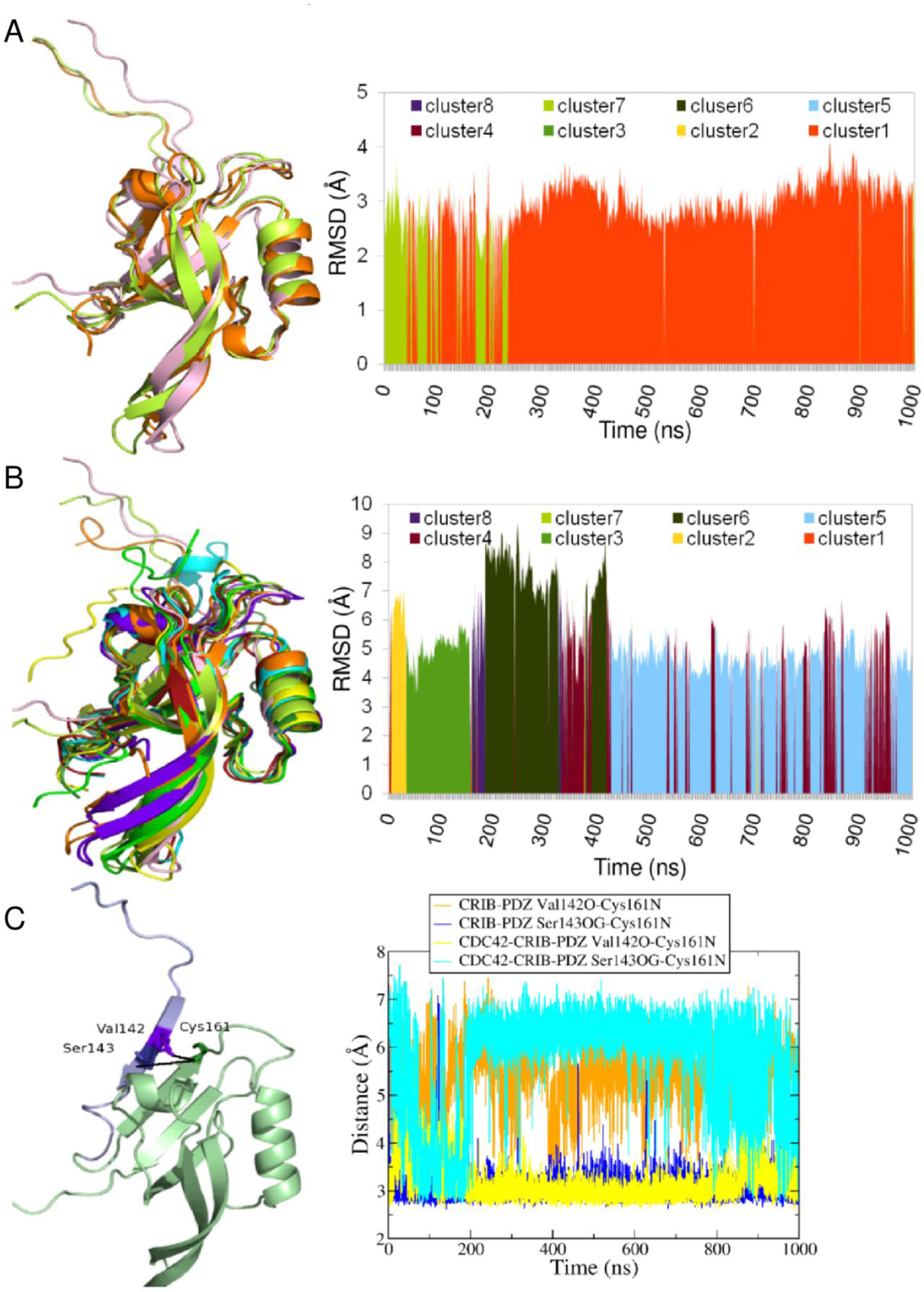
Altered conformations of Par6 PDZ in presence of CDC42 and CRIB.(A) and (B) are showing the cluster analysis of CRIB-PDZ with and without CDC42 simulations. Same colors are used in the RMSD plot and aligned structures. (C) Distance plot for Cys131N of PDZ, Val142O and Ser143OG of CRIB. A structure of CRIB-PDZ is also shown where PDZ is in pale green color and CRIB is in purple color. The residues involved in hydrogen bond are highlighted in dark colors as stick representation.

The hydrogen bonds between CRIB and PDZ in both the trajectories were analyzed which were stable for more than 60% of the simulation time. **Table S4** lists those interactions and it shows that there is only one differential interaction in presence or absence of CDC42 i.e., interaction of Cys161. When CDC42 is bound to CRIB, Cys161N of PDZ interacts with Val142O of CRIB and without CDC42, it interacts with Ser143OG **(Figure 9C)**. It seems that this shift in interaction leads to change in conformation from close to open carboxylate-binding loop and in other words in conveying the allosteric effect of CDC42. This interaction is very important in deciding the conformation of carboxylate-binding loop because substitution of corresponding residues in Drosophila Par6b PDZ to cysteine i.e., Q144C and L164C to facilitate disulfide bond formation leads to CRIB-PDZ which resembles CDC42 bound CRIB-PDZ in structure and function with significant increase in affinity for C-terminal ligands ^16^. When CDC42 interacts with CRIB, Asn39 from CDC42 forms hydrogen bond with backbone of Ser143 which moves it farther from PDZ ^30^. **Figure 10A and 10B** show this allosteric interaction network between the CDC42-CRIB-PDZ and CRIB-PDZ resulting in close or open state of carboxylate binding loop.

**Figure 10.**
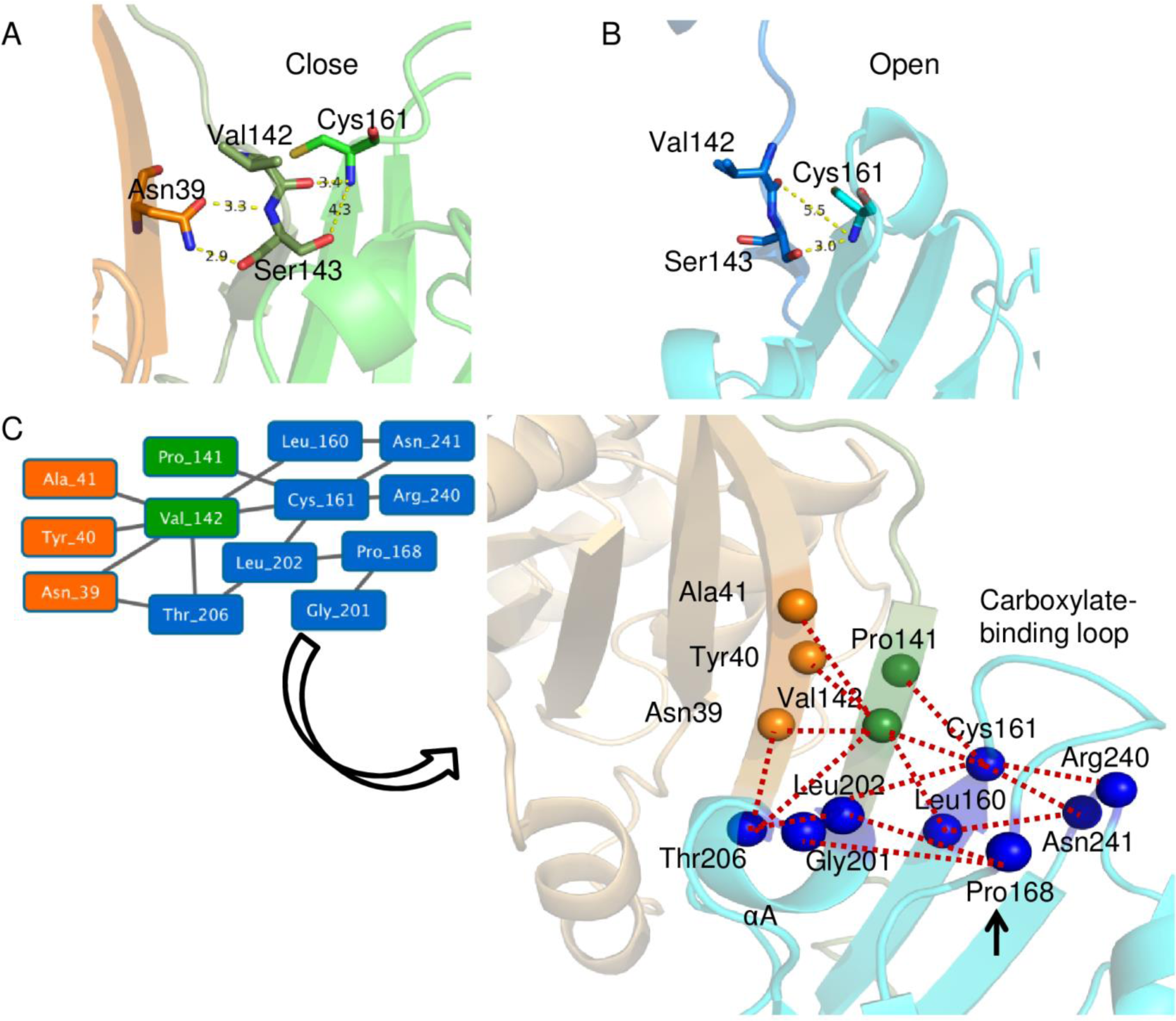
Allosteric residue interaction network of Par6 in presence and absence of CDC42.(A)Hydrogen bonds formed between Asn39 (CDC42) and Ser143 (CRIB) moved serine away from PDZ and hence Val142 (CRIB) interacts with Cys161 of carboxylate binding loop of PDZ leading to closed conformation of carboxylate binding loop in presence of CDC42. CDC42 is shown in orange color, CRIB is in dark green and PDZ in green color. (B) In absence of CDC42, Ser143 is hydrogen bonded to Cys161 and this results in open conformation of loop. CRIB and PDZ are shown in blue and cyan color respectively. (C) Dynamic residue interaction map created with Val142, Cys161, Pro168 and their first neighbors including interactions which are stable over 60% of the simulation time. Left panel network is generated with Cytoscape and the obtained residues were mapped on the structure of CDC42 (orange) and CRIB (green)-PDZ (cyan) complex to get the right panel figure. Residues forming dynamic interaction network are shown as Cα-spheres representing nodes and edges are shown in red color. Pro168 is marked with an arrow.

Though interactions between CRIB and PDZ in presence of CDC42 have been studied earlier using the crystal structure of CDC42 bound CRIB-PDZ ^30^ but the shift in hydrogen bonding pattern of Cys161 has never been discussed and related to its structural consequences. Since mutation of P171G (Drosophila numbering) which does not interact with CRIB or CDC42 directly leads to decoupling of effect of CDC42 on Par6 PDZ domain of Drosophila ^15^, we wanted to decipher the allosteric path followed for conveying the effect of CDC42 binding to other end of carboxylate binding loop. We created a dynamic intra-molecular interaction map for CDC42 bound CRIB-PDZ using distance cutoff 4.5Å between any two atoms including only those which are stable for more than 60% of simulation time (see methods). The important residues involved in hydrogen bonding identified from our simulation i.e., Cys161 and Val142 and Pro168 have been selected with their first neighbors in Cytoscape^31^ and this resulted in a residue interaction network showing interaction between Pro168 and other directly affected residues which is mediated by residues of αA helix of PDZ **(Figure 10C)**. Though our analysis shows the long range structural communications between Pro168 and residues of CDC42 and CRIB, but how this network is affected when Pro168 is mutated to glycine cannot be deduced with present data.

### Distance analysis of trajectories for close and open loop conformations

We have also used the distance of D169 (Cα atom) from the binding pocket of PDZ as a measure of the open and closed conformation (explained in methods section). This distance is 11.5 Å in the crystal structure of open conformation i.e., cognate internal complex and 9.3 Å in the crystal structure of closed conformation i.e., cognate C-terminal complex (**Figure S12A**). **FigureS12B** shows the distribution of the population of conformers in terms of the distance between Asp 169 and the peptide binding groove in cognate C-terminal complex varies between 7.1-13.5 Å and in cognate internal complex from 10.39 to 15.31 Å which peaks around 10.5 Å and 12.5 Å. These two distances correspond to two different conformations of carboxylate-binding loop namely open and close conformations. Both non-cognate complexes show dual peaks one for each open and close conformation. Very less proportion of close conformation in non-cognate internal complex is correlated with its faster transition. The distance distribution for both the ‘without peptide simulations’ showed shifting of population towards the closed conformation of loop whether it started from closed or open conformation of loop. In contrast, when distance distribution of CRIB-PDZ module is observed, it shows shift in population from close to open conformation and in presence of CDC42, it goes back to closed conformation which is more close and faster to unbound simulations **(Figure S12B)**. Other than this, the distribution plot confirms the importance of movement of D169 with respect to binding pocket as its distance from binding pocket can clearly define the open and closed state of carboxylate binding loop in the PDZ.

## Discussion

Penkert *et al*. ^10^ showed that when C-terminal peptide crystal structure is overlaid on the internal peptide Par-6 PDZ crystal structure, they show altered conformation of their carboxylate-binding loop with distances between backbone atoms from 3-7 Å. In this work, we have explored the mechanism of interaction for internal peptide and transition from open to close and close to open state of carboxylate binding loop. Explicit solvent MD simulations of microsecond time scale showed that the open and close conformation of loop are reversible and depends on the bound ligand. The time taken for conversion from one conformation to other is also ligand-dependent. Internal ligand is able to change the conformation of carboxylate-binding loop from close to open state in lesser time-scale with the help of Asp169 present in carboxylate-binding loop which pushes back the loop against the peptide Asp+1 residue due to repulsion. Though there are intramolecular PDZ interactions favoring close conformation of loop, overcoming those by repulsion and establishing strong interaction of Lys165 of PDZ and Asp+1 of peptide helps formation of open conformation. It seems that repulsion between aspartate residues is important for initial push of carboxylate binding loop and salt bridge provides additional energy for stability of open conformation. Since Par6 PDZ can readily change closed conformation of carboxylate binding loop to open conformation as illustrated by simulation on non-cognate internal complex, CDC42 has no effect on its internal peptide binding. This study provides the possible mechanism of internal peptide recognition and the conservation profile of important residues in other similar internal peptide binding PDZ domains highlights that proposed mechanism might be extendable to other similar PDZ domains also. Thus we are able to predict 47 PDZ domains to be capable of interacting internal peptide from human PDZome. The simulations on CASK PDZ, DLG1-1 PDZ and MAST1 PDZ further validated our observations. A recently published article indicated that Pals1 PDZ can interact with internal ligands based on its structural resemblance to Par6 PDZ internal peptide complex ^32^ and it also possess critical residues involved in internal peptide interaction as proposed by our simulation results.

MM-PBSA analysis is in agreement to the experimental dissociation constant data that without CDC42, C-terminal peptide has low affinity for PDZ domain. Thus, explaining the requirement of CDC42 for forming high affinity C-terminal complex ^15^. Non-cognate and cognate internal complex have similar MM-PBSA energies illustrating independence of internal peptide interaction from CDC42 as suggested in the literature ^10^. This confirms that MM-PBSA may readily be used for differentiating C-terminal versus internal peptide interactions of PDZ domain.

MD simulations on CRIB-PDZ with and without CDC42 showed change in conformation of carboxylate-binding loop from close to open with shift in interaction of Cys161-Val142 to Cys161-Ser143 and thus switching preference of C-terminal peptide recognition to internal peptide interactions. This same Ser143 of CRIB interacts with CDC42 which makes it unavailable for PDZ in CDC42 bound complex. Without CRIB/CDC42, Par-6 PDZ alone has propensity to form closed conformation in absence of any peptide which highlights its preference for dominant mode of closed conformation without any assistance of external factors. We show that other proteins and adjacent domains can modulate the PDZ interactions thus results in a regulated complex assembly by PDZ domains.

## Methods

### Compilation of starting structures for MD simulations

The crystal structures of C-terminal peptide ligand bound Par-6 PDZ domain (PDB ID: 1RZX) ^15^ and internal peptide bound Par-6 PDZ domain (PDB ID: 1X8S ^10^) were downloaded from PDB (http://www.rcsb.org). These two complexes are referred as cognate C-terminal and cognate internal peptide complex respectively. The non-cognate C-terminal peptide complex was generated by removing the native internal peptide coordinates from cognate internal peptide complex (1X8S) and then transforming the coordinates of the C-terminal peptide from cognate C-terminal peptide complex (1RZX) into the binding pocket of 1X8S after optimum superposition of the coordinates of the PDZ domains alone. Similarly the non-cognate internal peptide complex was generated by removing the native C-terminal peptide coordinates from cognate C-terminal peptide complex (1RZX) and then transforming the coordinates of the internal peptide from cognate internal peptide complex (1X8S) into the binding pocket of 1RZX. The peptide free unbound structures for Par-6 PDZ domain were also generated by removing peptide coordinates from 1X8S and 1RZX, and used as starting structures for simulations on ligand free Par-6 PDZ. The crystal structure of Cdc42 bound CRIB-PDZ domain (PDB ID: 1NF3) ^30^ was obtained from PDB and used as starting structure for simulations on effector bound ligand free PDZ domain. The starting structure of CRIB-PDZ in absence of effector was obtained from 1NF3 by removing coordinates of Cdc42. The PDB entries 1KWA ^33^, 3RL7 ^34^ and 3PS4 were used as starting structures for MD simulations on CASK, DLG1-1 and MAST1 PDZ domains.

### MD simulations

The explicit solvent MD simulations were carried out by AMBER 12 ^35^ package using ff03 force field ^36^. The structures were solvated using TIP3P ^37^ water model such that rectangular solvent boxes extended of 8 Å away from the outer most protein atoms in X, Y and Z directions. The net charge of the solvated system was then neutralized by addition of required number sodium or chloride counter ions. All the systems were simulated using these Periodic boundary conditions. The long-range electrostatic interactions were computed in reciprocal space using particle-mesh Ewald method (PME) ^38^ and a cut off of 10Å was used for non-bonded interactions in direct space. Langevin temperature equilibration scheme was used to maintain the temperature of the system. Constant pressure periodic boundary of an average pressure of 1 atm was used with isotropic position scaling to maintain the pressure. The various solvated systems were minimized using steepest descent algorithm initially and then conjugate gradient approach to an RMS gradient of 0.001 kcal/mol/Å. After minimization, they were allowed to heat up gradually from 0 K to 300 K over 20 ps of MD simulation at constant volume. The second equilibration step consisted of 100ps of MD simulation under NPT conditions. After equilibration production MD simulations of 500ns or 1 μs were performed under NPT conditions for each of the systems using time step of 2 fs. The bonds involving hydrogen were constrained using SHAKE alogorithm. Snapshots were saved at an interval of 100 ps for subsequent analysis. The simulations were carried out on workstations with NVIDIA GPUs using PMEMD (particle mesh Ewald molecular dynamics) module of AMBER. The MD trajectories were analyzed using PTRAJ module of AMBER 12 and in-house perl scripts.

### Analysis of hydrogen bonds in MD trajectories

The analyses of various types of hydrogen bonds in structures sampled during the MD simulation were carried by using hbond command of the cpptraj module of AMBER package. The criteria for defining hydrogen bonded atoms pairs involved a donor-acceptor distance of less or equal to 3.5Å and donor-hydrogen-acceptor angle of 135° or higher. The hydrogen bonds which remained intact for more than 60% of the simulation length were considered as dynamically stable hydrogen bonds.

### Cluster analysis of MD trajectories

The snapshots from MD trajectories of 1 μs length were extracted at an interval of 1 ns using PTRAJ module of AMBER. This resulted in 1000 conformers per simulation. The conformers from cognate and non-cognate C-terminal and internal peptide complex simulations (total 4000 conformers) were clustered together using the kClust utility of the MMTSB (Multiscale Modeling Tools for Structural Biology) suite ^39^ on the basis of their Cα RMSDs. The clustering radius of 3Å was used. The second set of clustering involved snapshots from trajectories cognate complexes with snapshots from MD trajectories of unbound structures of C-terminal and internal peptide specific PDZ domains. The second clustering was also based on Cα RMSDs of PDZ domain alone and used a clustering radius of 3Å. The third set of clustering was also done for the MD trajectories of Cdc42 bound CRIB-PDZ and effector free CRIB-PDZ module on the basis of the Cα RMSDs of CRIB-PDZ only. Since CRIB domain is disordered in absence of Cdc42, higher clustering radius of 4 Å was used for CRIB-PDZ. The percentage occupancy or population of conformers in each cluster was calculated by counting the fraction of conformers present in each cluster out of the total number of conformers in the trajectory for each simulation.

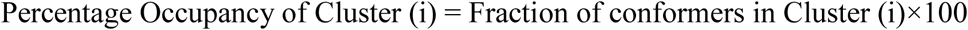

### Analysis of open & closed conformations of PDZ domain

The carboxylate-binding loop moves close to or far away from the Par-6 PDZ binding pocket during the transition from open to close state or vice versa. Therefore, the distance between D169 of Par-6 PDZ carboxylate-binding loop and its binding pocket was used as the criteria for distinguishing between open and closed states. Since G173 and V239 residues in the binding pocket showed much less movement during the simulations and remained essentially fixed, distance between Cα atom of D169 and the mid-point of the Cα atoms of G173 and V239 was used to define closed and open states of PDZ domain. The corresponding residues in CDC42:CRIB-PDZ (PDB ID: 1NF3) were E166(D169), G170(G173) and I236(V239).

### MM-PBSA analysis

In order to calculate the binding free energies for cognate and non-cognate peptides in C-terminal and internal peptide complexes, 2000 frames at 500 ps interval were retrieved from each of the four 1 μs trajectories for various cognate and non-cognate complexes of Par-6 PDZ domain. The solvent molecules were removed from the snapshots and implicit solvent MM-PB/SA calculations were performed on these trajectories using python module of MM-PB/SA package of AMBER 12 ^40^. The binding free energy for the peptide was calculated by subtracting the free energy of peptide and PDZ domain from free energy of the whole complex. For MM-PB/SA calculations on CASK, MAST1 and DLG1-1 PDZ-peptide complexes, 1000 frames were extracted at 500 ps interval because the simulation lengths were 500ns only.

### Multiple sequence alignment of PDZ domains

Sequences of internal peptide binding PDZ domains were obtained from the published work of Mu *et al*. ^7^ and multiple sequence alignments were carried out using Clustal Omega software ^41^. All human PDZ sequences were taken from te velthius *et al.*^4^ and only which could be mapped to Uniprot ID ^42^ were used for alignment. Alignment was visualized with Jalview^43^.

### Generating dynamic residue interaction network

The distance between all atoms of Cdc42 bound CRIB-PDZ complex (inter/intra-residue with intermediate hydrogen atom) is obtained using cpptraj hbond command with their occupancy in frames extracted at 100ps interval. Using distance cutoff of 4.5Å and occupancy of more than 60%, a filtered interaction map of residues (whose any atom is in contact) is obtained where interaction between ±2 neighboring residues is ignored to get long range structural communications. This complex residue interaction map is loaded in Cytoscape^31^ and important residues identified during simulation (Val142 and Cys161) and Pro168 is selected with their first neighbors to get a new interaction network which is focused on these important residues. The resulted network had 13 nodes and 15 edges.

## Author Contributions

N.S. performed the experiments. N.S. and D.M. planned the experiments and wrote the paper.

## Acknowledgment

This work was supported by the Department of Biotechnology, Government of India grant to National Institute of Immunology, New Delhi. DM also acknowledges financial support from Department of Biotechnology, India under BTIS project (BT/BI/03/009/2002) and COE grant (BT/COE/34/SP15138/2015). NS was supported by fellowship from CSIR, India.

## Supporting Information

Supplementary tables and figures (PDF)

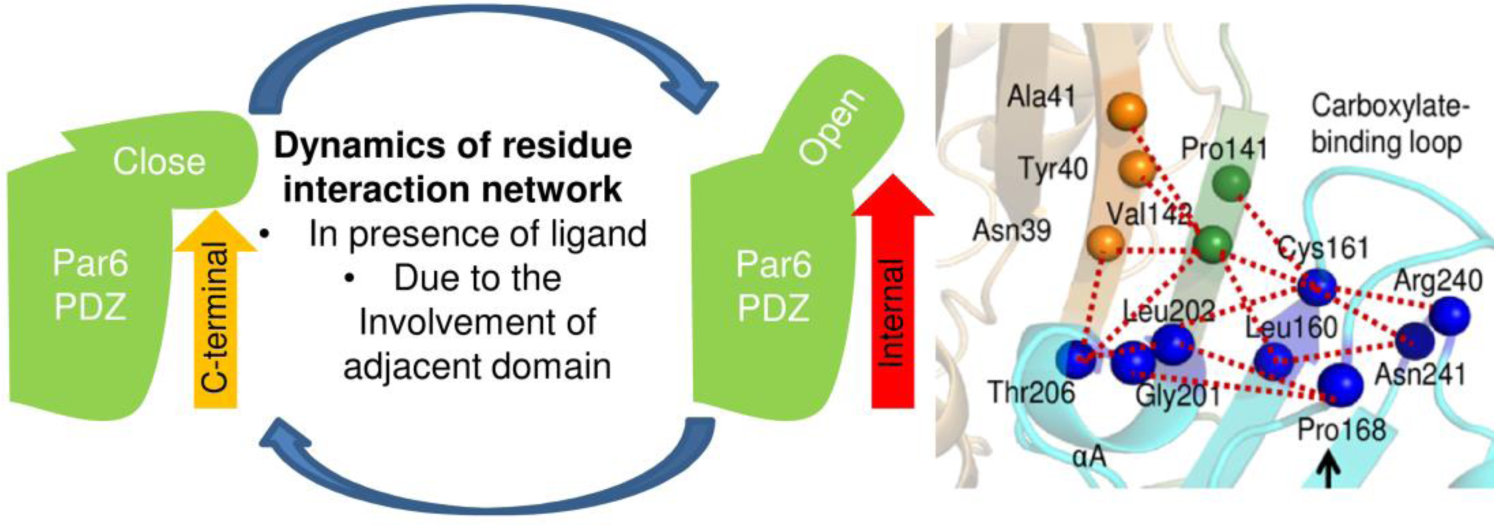

