## Supplementary file for "Structural basis of internal peptide recognition by PDZ domains using molecular dynamics simulations"

### Supplementary Tables and Figures

**Table S1.** Percentage occupancy of the clusters formed during clustering of cognate C-terminal, internal and both the without peptide simulations of 1000ns.

| Cluster | C-terminal complex | C-terminal WOP | Internal complex | Internal WOP |
|---|---|---|---|---|
| 1 | 13.8 | 86 | 0.1 | 0.1 |
| 2 | 0.8 | 0 | 0 | 64.8 |
| 3 | 0.2 | 0.6 | 99.3 | 4.7 |
| 4 | 50.8 | 0 | 0 | 0 |
| 5 | 32.6 | 8.2 | 0 | 17.5 |
| 6 | 1.8 | 5.2 | 0.6 | 12.9 |

**Table S2.** Distance between D169 of carboxylate-binding loop and D(+1) of internal peptide/ V(0) of C-terminal peptide. Carboxylate group's oxygen atoms were used for distance calculation.

| Structure | PDZ Residue | Peptide Residue | Shortest distance (Å) |
|---|---|---|---|
| Cognate internal Complex | D169 OD1/OD2 | D(+1) OD1/OD2 | 9.9 |
| Non-cognate internal Complex | D169 OD1/OD2 | D(+1) OD1/OD2 | 5.9 |
| Cognate C-terminal Complex | D169 OD1/OD2 | V(0) O/OXT | 9.8 |
| Non-cognate C-terminal Complex | D169 OD1/OD2 | V(0) O/OXT | 14.0 |

**Table S3.** Percentage occupancy of the clusters formed during clustering of CRIB-PDZ module simulated in presence and absence of CDC42.

| Cluster | CDC42-CRIB-PDZ | CRIB-PDZ |
|---|---|---|
| 1 | 85.4 | 0.3 |
| 2 | 0 | 3.1 |
| 3 | 0 | 12.3 |
| 4 | 0 | 17.3 |
| 5 | 0 | 45.7 |
| 6 | 0 | 17.6 |
| 7 | 14.6 | 0.2 |
| 8 | 0 | 3.5 |

**Table S4.** Hydrogen bonds found between CRIB and PDZ domain in presence and absence of CDC42 stable over 60% of the trajectory of 1000ns.

| S.No. | Hydrogen bond between CRIB-PDZ with CDC42 (% occupancy) | Hydrogen bond between CRIB-PDZ without CDC42 (% occupancy) |
|---|---|---|
| 1 | **S143OG-C161N (91.38)** | **V142O-C161N (70.44)** |
| 2 | S144O-R159N (98.71) | S144O-R159N (98.76) |
| 3 | S144N-R159O (85.32) | S144N-R159O (92.81) |
| 4 | I146N- R157O (93.03) | I146N-R157O (99.89) |

**Supplementary Figures**

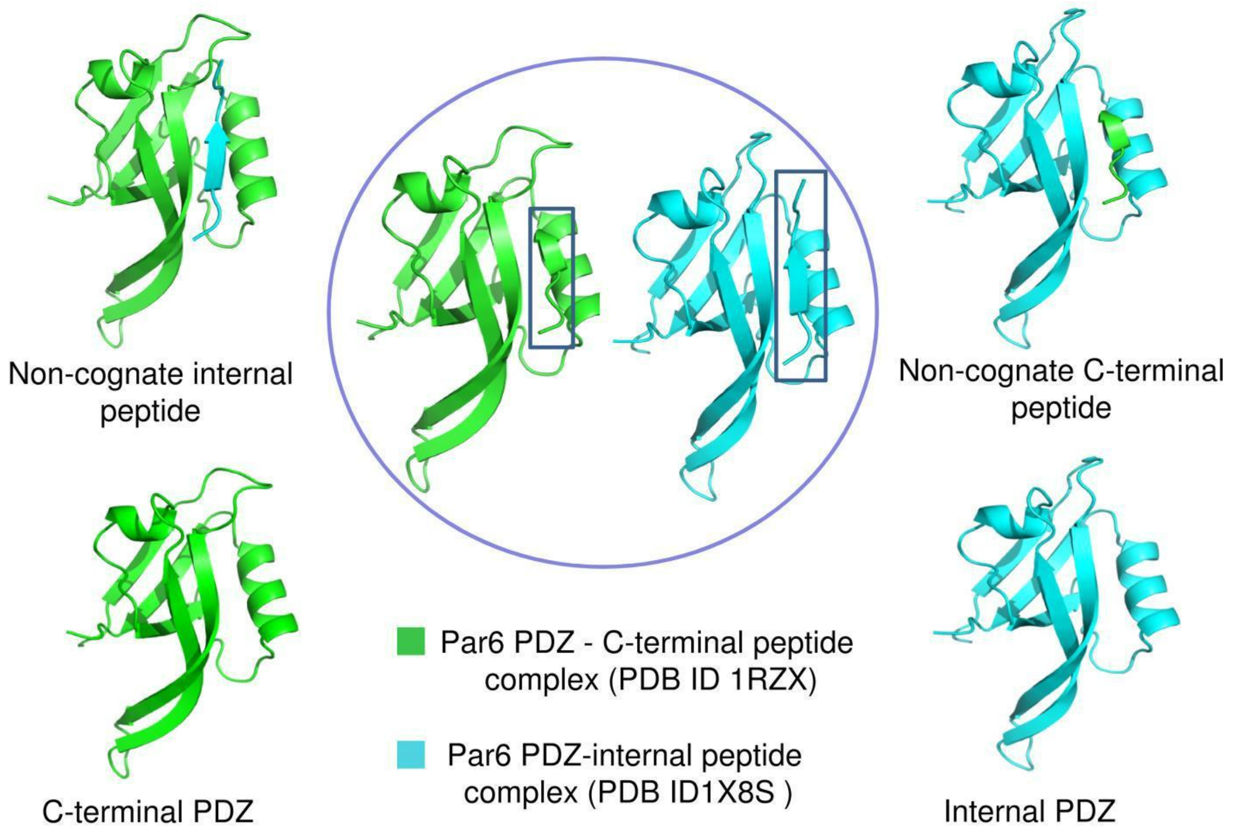


**Figure S1.** Cognate and non-cognate structures of Par6 PDZ used for MD simulations. C-terminal and internal peptide complexes are shown in center (encircled in blue color). The structures generated after interchanging their peptides (non-cognate complexes) and removal of peptides are also shown.

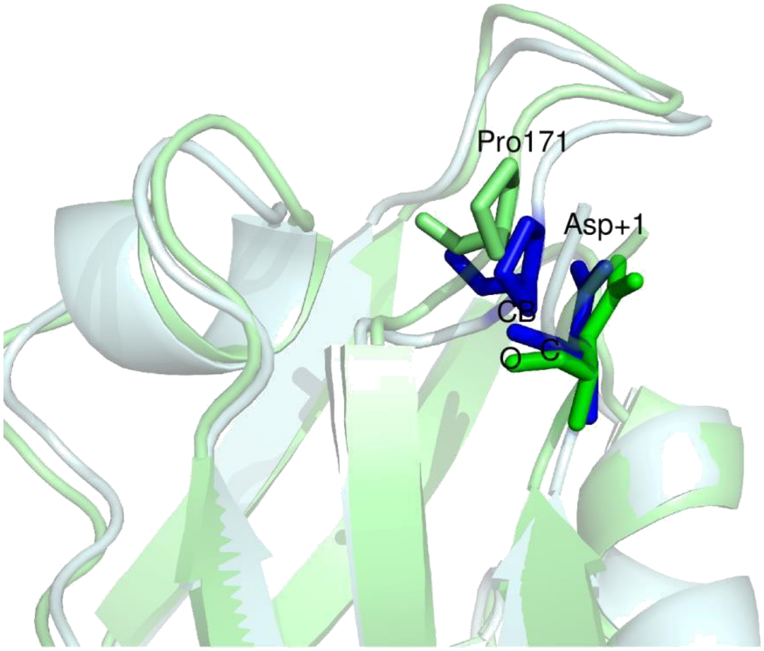


**Figure S2.** Close contacts between carboxylate-binding loop of PDZ and internal peptide in non-cognate internal complex. Non-cognate internal complex in pale cyan color, is showing steric clashes between CB of Pro 16 and C and O of Asp (+1) highlighted in blue color. Close contacts were computed by Xleap package of AMBER 12. After minimization steric clashes were relieved by small movement in carboxylate-binding loop and peptide shown in pale green color with the side chains of Pro 16 and Asp (+1) represented in green color.

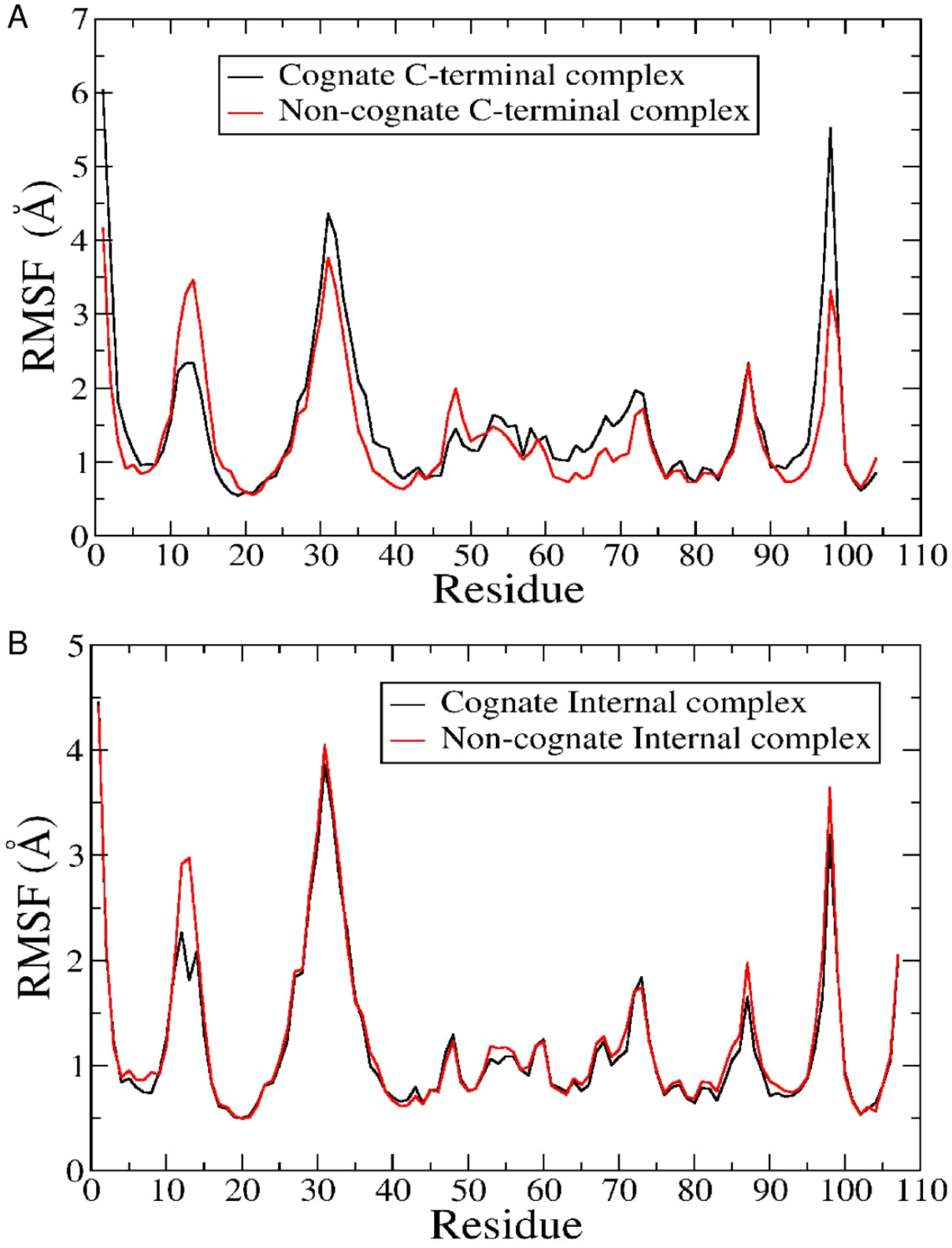


**Figure S3.** RMSF plots for peptide bound simulations. (A) RMSF plots for non-cognate and cognate C-terminal complex and (B) RMSF plot for non-cognate and cognate internal complex are indicating high fluctuations in carboxylate-binding loop (residues 9-16).

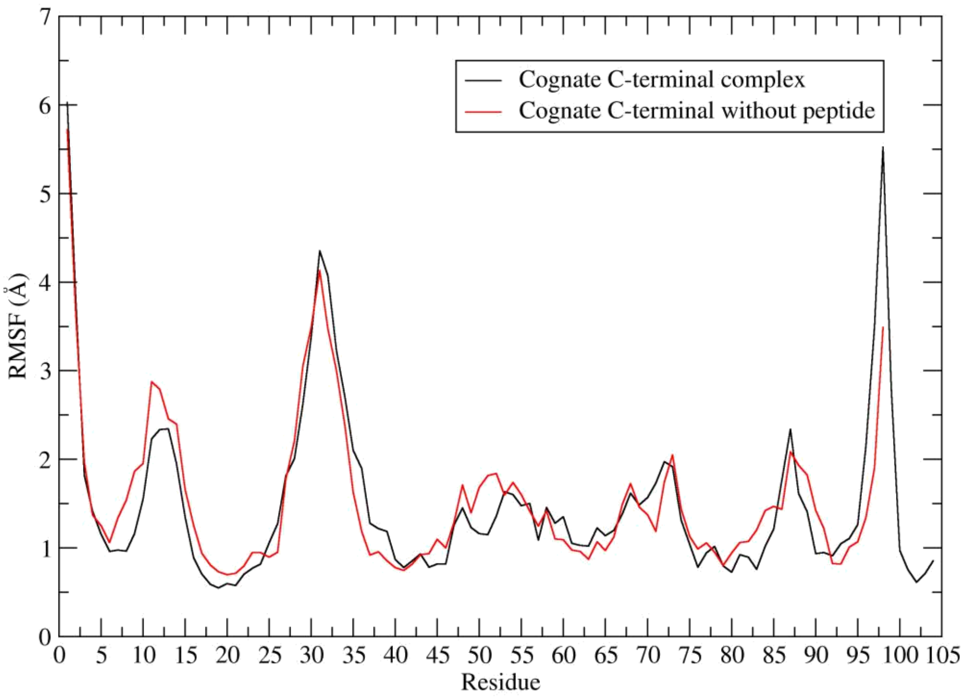


**Figure S4.** RMSF plot for cognate C-terminal and unbound C-terminal complex. Residues 9-16 represent carboxylate-binding loop with change in conformation

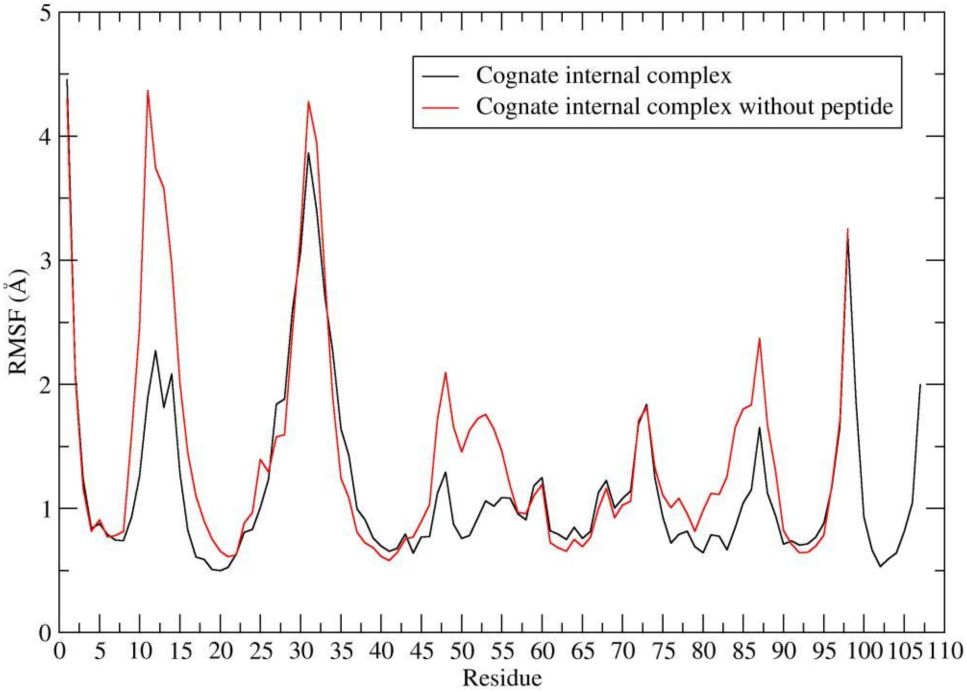


**Figure S5.** RMSF plot for cognate internal and unbound internal complex. Residues 9-16 represent carboxylate-binding loop with change in conformation

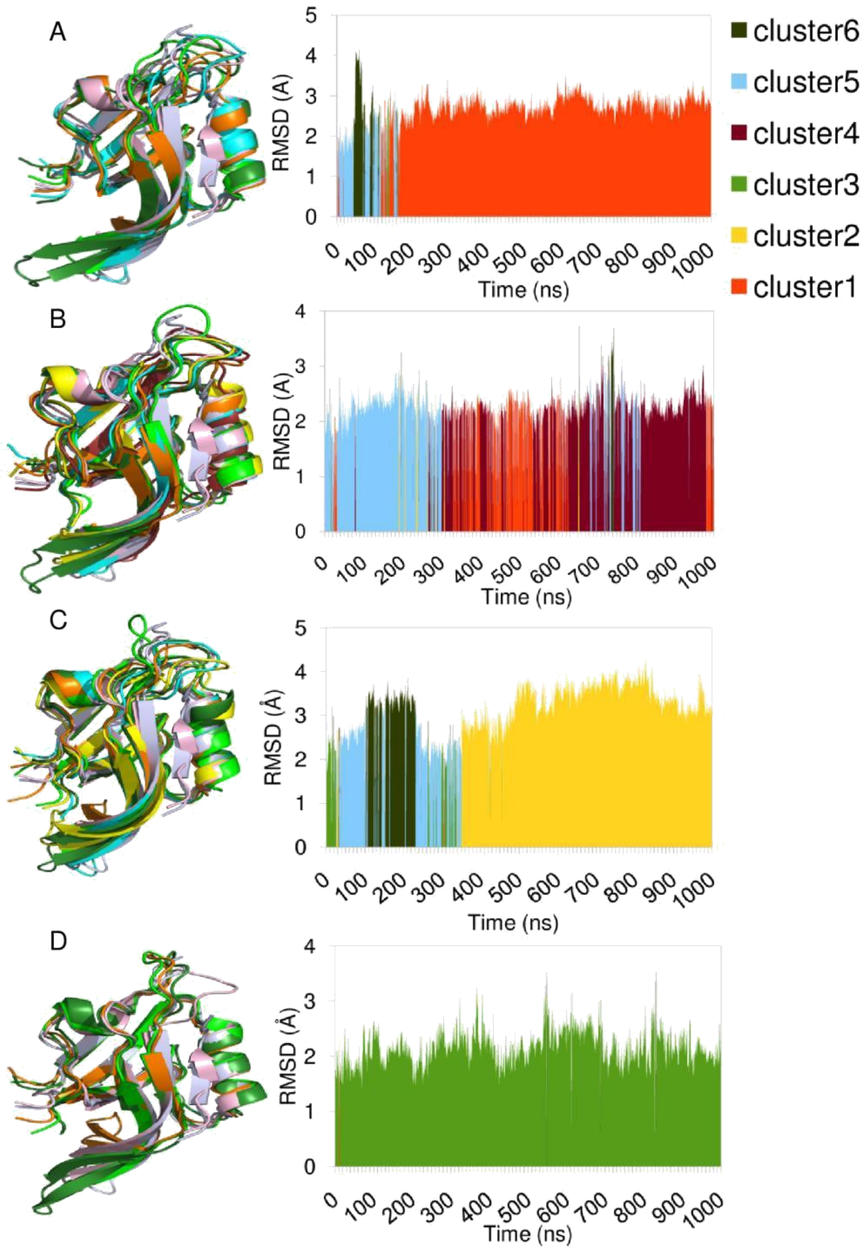


**Figure S6.** Cluster analysis of cognate and 'without peptide' C-terminal and internal complex trajectories. (A) Cognate C-terminal without peptide (B) Cognate C-terminal complex (C) Cognate internal without peptide and (D) Cognate internal complex. Each of the six clusters is shown in different colors which are followed in all of these four simulations and the color used for the representative member is same as the cluster color. C-terminal and internal complex crystal structures are shown for reference in the same color as in Figure 6.

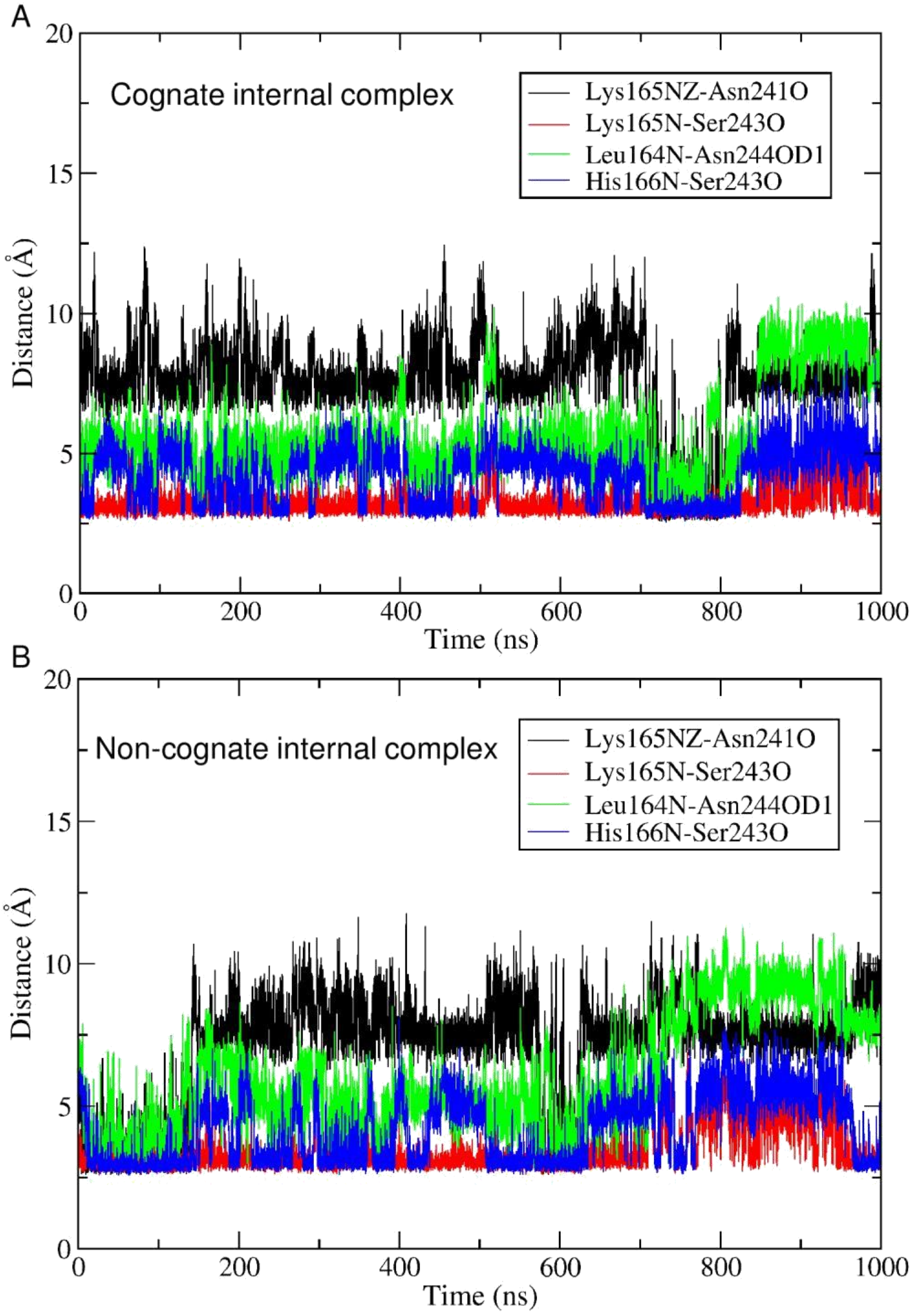


**Figure S7.** Distance *vs* time plot. Interactions between carboxylate binding loop and αB-βF loop identified at the intermediate stage of non-cognate internal complex were measured for (A) cognate internal peptide complex and (B) non-cognate internal peptide complex. Four interactions are shown in four different colors.

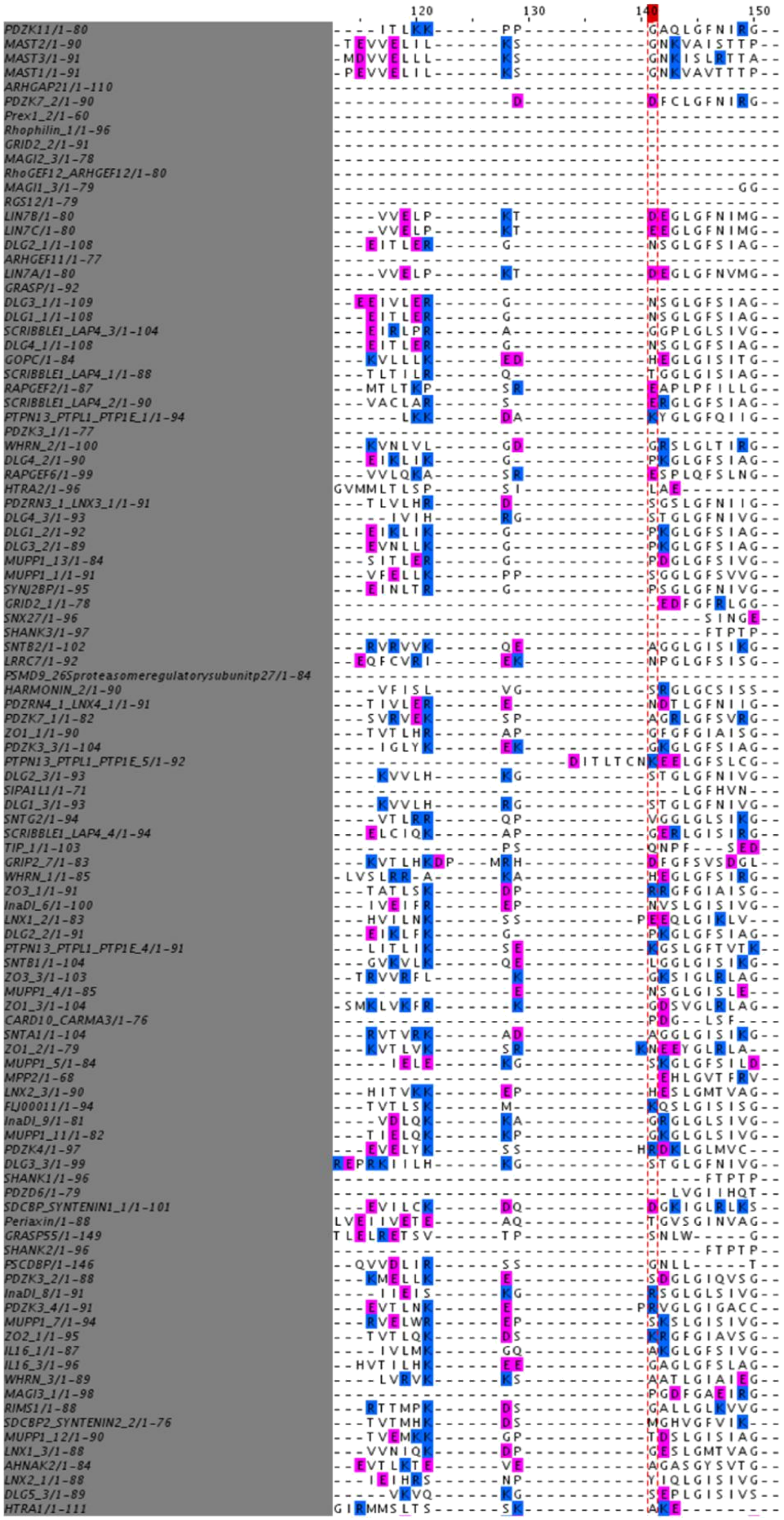

**Figure S8a (Cntd.)**

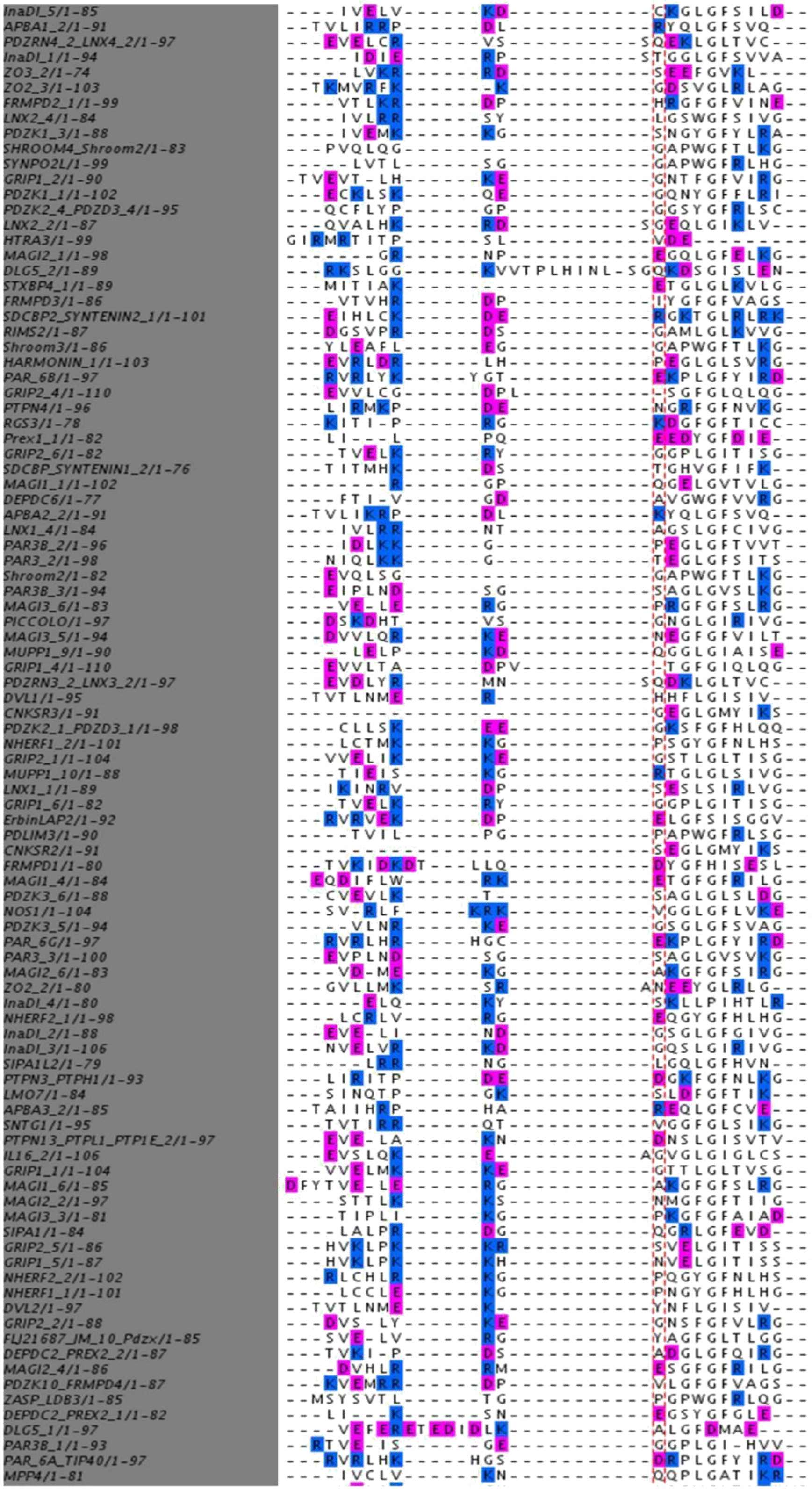

**Figure S8b (Cntd).**

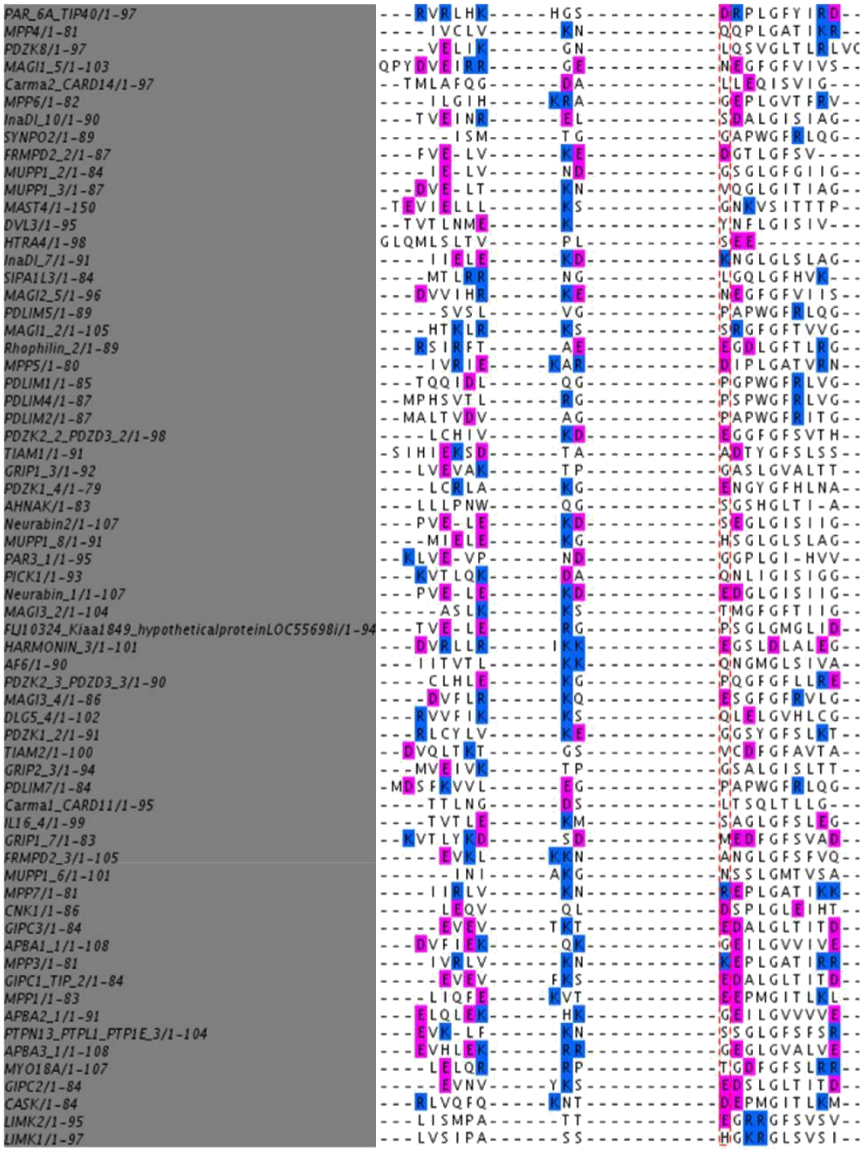

**Figure S8c**. MSA of human PDZome. 264 human PDZ domains were aligned using Clustal Omega and corresponding D169 position is marked in dashed red color box.

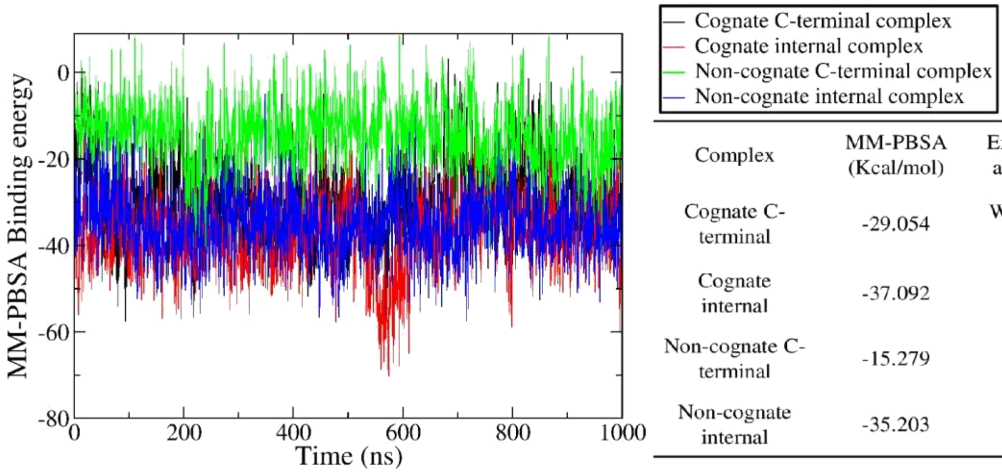


| Complex | MM-PBSA (Kcal/mol) | Experimental $K_D$ in μM as reported in literature |
|---|---|---|
| Cognate C-terminal | -29.054 | Without CDC42: 72 ± 5<br>With CDC42: 6 ± 2 |
| Cognate internal | -37.092 | Without CDC42: 8<br>With CDC42: 6 |
| Non-cognate C-terminal | -15.279 | - |
| Non-cognate internal | -35.203 | - |

**Figure S9.** MM-PBSA analysis is in accordance of experimental dissociations constant values. Left panel is the MM-PBSA energy plot for 1000 ns for all four simulations of complexes and average MM-PBSA binding energies for last 500ns are shown at right with the experimentally measured dissociation constants values taken from literature.

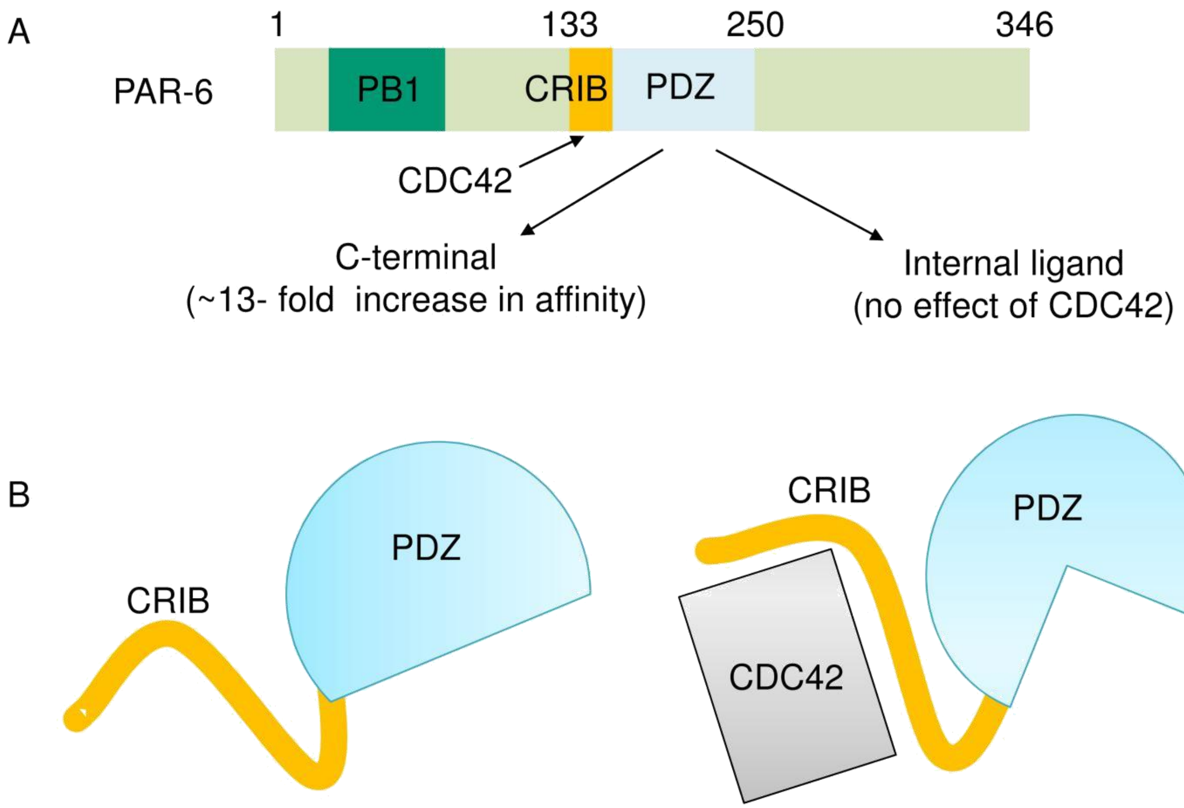


**Figure S10.** (A) Domain organization of Par6 protein. CRIB domain is present adjacent to PDZ domain. (B) CDC42 interacts with CRIB. Binding of CDC42 to CRIB allosterically impact binding of C-terminal peptides to PDZ and results in 13-fold increase in affinity for C-terminal peptides.

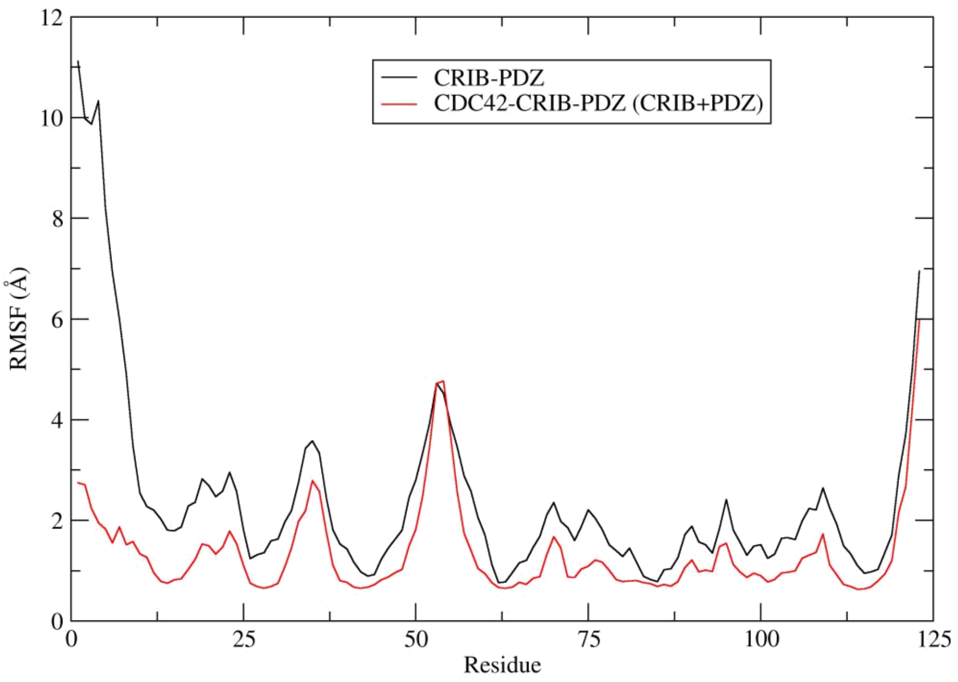


**Figure S11.** RMSF plot for CDC42 bound CRIB-PDZ and CRIB-PDZ.

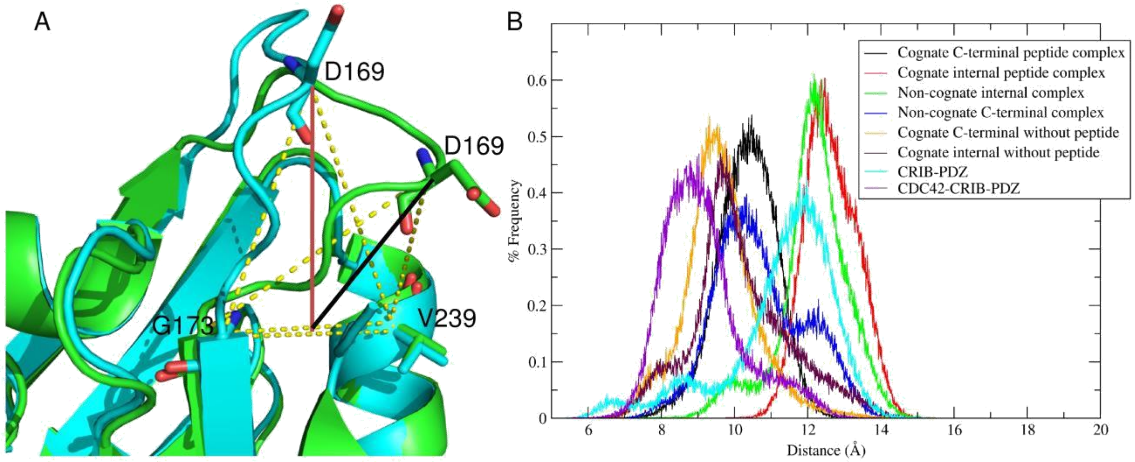


**Figure S12.** Distribution of the open and closed conformations in all six MD runs defined by the distance between the D169 and peptide binding pocket.(A), Red line represents the distance in cognate internal complex (11.5 Å) and black line is for cognate C-terminal complex (9.3 Å) crystal structure. (B) Distribution of distance computed over entire trajectories showing cognate C-terminal (black) and internal (red) complex sample significantly different conformations during simulation having peaks at around 10.5 Å and 12.5 Å respectively. Non-cognate complexes (green and blue) show two peaks indicating mixed populations of open and closed conformation. Without peptide simulations (orange and violet) have peaks at less than 10.5 Å indicating their closed conformation of carboxylate-binding loop. With CDC42 and without any peptide, carboxylate-binding loop of PDZ remains in closed state while after removing CDC42, it goes to open conformation.
